# The long noncoding RNA *Dory* is required for female but not male spatial learning and memory

**DOI:** 10.64898/2026.04.01.715990

**Authors:** Seonghee Jung, Mitchell J. Cummins, Saba Altaf, Rowan J. Heggen, Anne Poljak, Lachlan Ferguson, Kelly Clemens, Fabien Delerue, Lars M. Ittner, John S. Mattick

## Abstract

Large numbers of long noncoding RNAs (lncRNAs) exhibit region- or cell-specific expression and subcellular locations in the mammalian brain. We analyzed the expression and function of the mouse lncRNA *2700046G09Rik* (also called *Sgms1os1*), which we have named *Dory* due to sex-specific disruption of spatial memory in rodent gene knockout and RNA knockdown. We show that *Dory* is predominantly expressed in a punctate pattern in nuclei of excitatory neurons in the hippocampus, and in both neuronal and non-neuronal cells in the cerebellum. We show by high resolution RNA sequencing that *Dory* brain transcripts are composed of 1-3 exons with multiple isoforms. Deletion of the constitutive first exon of *Dory* by CRISPR-Cas9 genome editing resulted in the impairment of spatial memory and some aspects of balance in female but not male mice. *Dory*-null mice presented without overt changes in brain morphology. Knockdown of the rat homolog of *Dory* in dorsal hippocampus using antisense oligonucleotides confirmed inhibition of spatial memory in females only. RNA sequencing and mass spectrometry revealed differential hippocampal gene and protein expression profiles, notably of prolactin, growth hormone and pro-opiomelanocortin, between male and female *Dory* knockout mice.

## Introduction

Long noncoding RNAs (lncRNAs), defined as intergenic, intronic or antisense transcripts greater than 500 nt in length that have little or no protein-coding potential, are pervasively expressed from the mammalian genome [1]. Many lncRNAs are highly alternatively spliced [2] and many, if not most, appear to be expressed from genetic loci called enhancers [1, 3], which control the spatiotemporal patterns of gene expression.

A large fraction of mammalian lncRNAs are expressed in highly specific patterns in the brain [4–9], with evidence for the participation of lncRNAs in brain development [10–18], neurogenesis and neuronal differentiation [19–30], synaptic function [31–38], learning and memory [39–43], cognition [30], locomotor control [44] and behavior [45]. LncRNAs have also been implicated in stress, neurological diseases and psychiatric conditions [10, 12, 14, 28, 46–50]. Many brain-expressed lncRNAs show evolutionary conservation among vertebrates [51], whereas others are primate- or human-specific [5, 52]. Recent studies have shown that modification of a synaptic lncRNA is required for fear extinction [53].

However, little is known about the function of the majority of lncRNAs that are expressed in the brain and, indeed, most have yet to be identified. The lncRNA *2700046G09Rik*, which was first identified by high throughput RNA sequencing as part of the RIKEN FANTOM project [54, 55], and is also known as *Sgms1os1* [56] and we now term *Dory*, was shown by the Allen Brain Atlas (ABA) using *in situ* hybridization (ISH) to be highly expressed in the hippocampus and cerebellum (Fig. 1) [4, 57]. Due to this spatial distribution, we hypothesized that *Dory* may have a role in hippocampus-dependent memory and cerebellum dependent motor control.

**Fig. 1:**
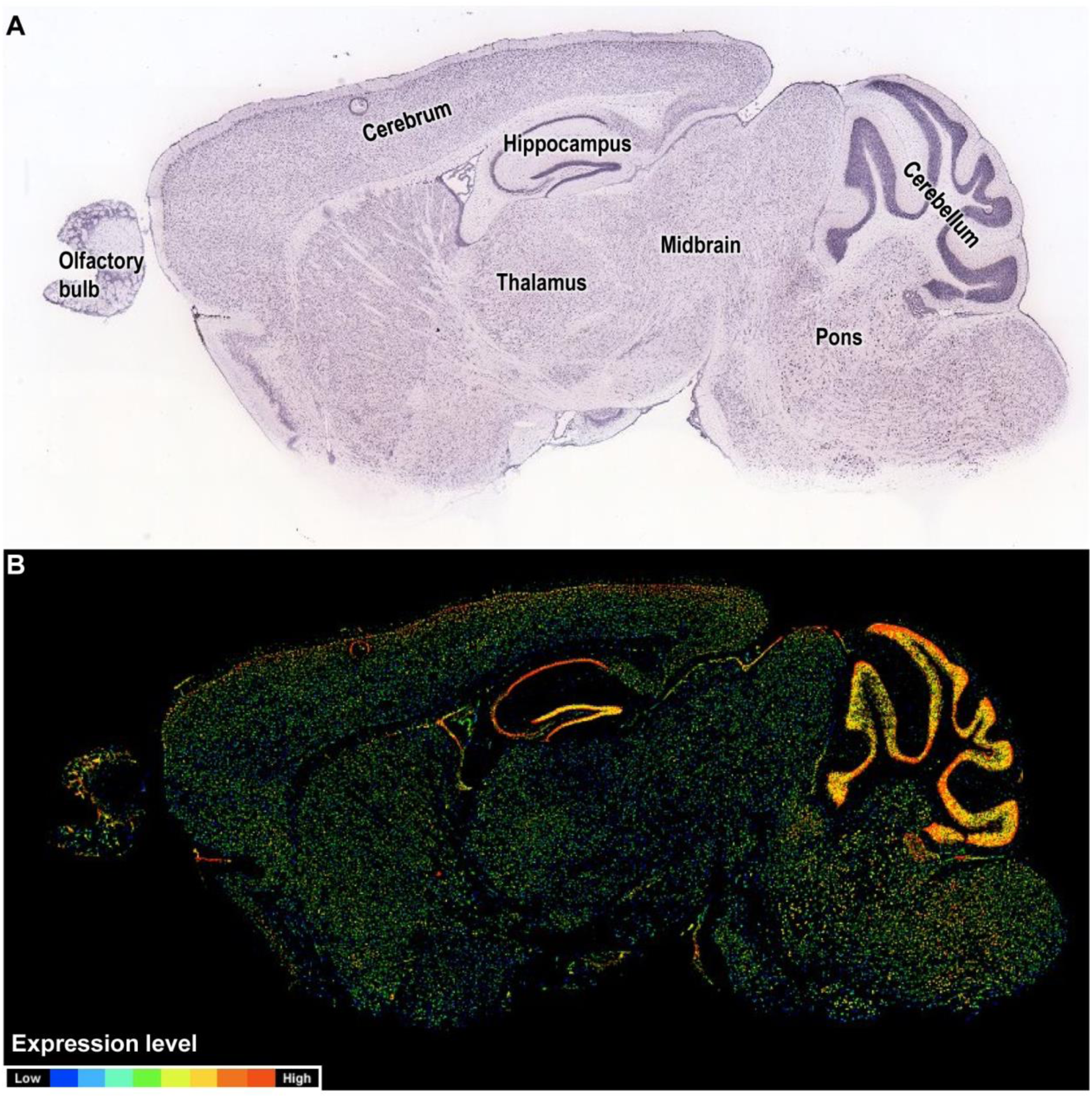
*In situ* hybridization (ISH) of sagittal sections with *Dory* (*2700046G09Rik*/*Sgms1os1*). Images were obtained from the Mouse Brain Atlas (http://mouse.brain-map.org/gene/show/43031) of the Allen Institute for Brain Science [57]. **(A)** Primary ISH hybridization data **(B)** Computer transformation whereby the average ISH signal intensity within a connected area is color-coded to range from blue (low expression intensity), through green (medium intensity) to red (high intensity).

We confirmed the presence of *Dory* in mouse hippocampus and cerebellum, and found enrichment in *Rbfox3*^+ve^ nuclei in all hippocampal subregions but not cerebellar nuclei. We then generated brain transcript models for *Dory* by high resolution RNA capture sequencing and used CRISPR-Cas9 genome editing to delete the constitutive first exon, followed by behavioral and molecular phenotyping in wild-type and *Dory* knockout mice. We found that knockout mice had deficits in spatial memory and some motor skills in females but not males, and named this lncRNA *Dory* after the amnestic female fish from the movie *Finding Nemo*. We also identified molecular factors that may contribute to these differences. We confirmed that direct knockdown of *Dory* in rat dorsal hippocampus results in sex-specific impediment of spatial memory.

## Results

### *Dory* is expressed in the nucleus of mature neurons in the hippocampus

The ABA showed that *Dory* (*2700046G09Rik*/*Sgms1os1*) is expressed in the *cornu ammonis* (CA) and the dentate gyrus (DG) of the hippocampus as well as in the cerebellum in mouse brains (Fig. 1), but knowledge of its spatial expression in different cell types and brain regions is limited. To supplement the ISH evidence from the ABA, we performed RNAscope targeting *Dory* in adult wild-type mouse brain sections (Fig. 2). Analysis of hippocampal subregions confirmed expression of *Dory* in CA subregions CA1 to CA3 and in the DG. Notably, *Dory* RNAscope signals were more intense in CA2 and DG than CA1 and CA3, and localized in punctate subnuclear domains of *Rbfox3*^+ve^ hippocampal neurons, with little detected in *Rbfox3*^−ve^ non-neuronal nuclei (Fig. 2A, B). *Dory* was expressed in both excitatory (*Grin1*^+ve^) and inhibitory (*Gabrg1*^+ve^) neurons in the hippocampus (Supplementary Fig. S1), and in immature neurons (*Dcx*^+ve^) and neural stem cells (*Nestin*^+ve^) in the DG (Supplementary Fig. S2). In the cerebellum (CB) there was approximately equal expression of Dory in *Rbfox3*^+ve^ neuronal nuclei and *Rbfox3*^−ve^ non-neuronal nuclei (Fig. 2A, B) in all layers including *Pcp2*^+ve^ Purkinje cells (Supplementary Fig. S2). *Dory* was also expressed in *Chat*^+ve^ neurons sparsely throughout the brain (Supplementary Fig. S2). Our data indicates that *Dory* is mainly present in neuronal nuclei in the hippocampus and present in both neuronal and non-neuronal nuclei in the cerebellum. There was no discernable difference in the expression patterns between male and female mice (Fig. 2C).

**Fig. 2:**
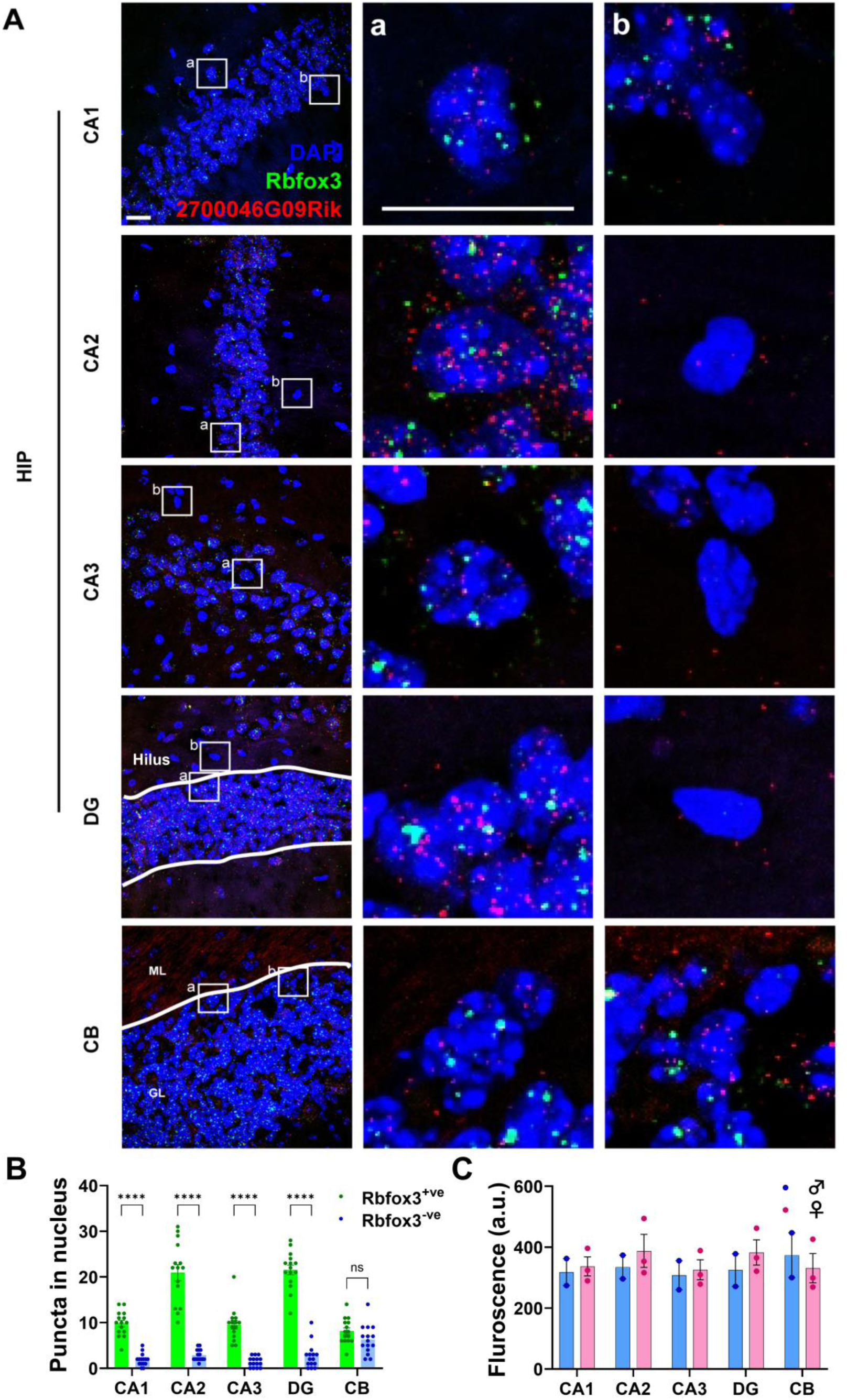
RNAscope *in situ* hybridization for *Dory* (*2700046G09Rik/Sgms1os1*)^+ve^ in hippocampal subregions CA1, CA2, CA3 and DG; and cerebellum (CB). **(A)** Representative image of (a) *Dory*^+ve^ and *Rbfox3*^+ve^ nuclei and (b) *Dory*^+ve^ and *Rbfox3*-ve nuclei. **(B)** Quantification of *Dory* puncta in the nucleus of *Rbfox3*^+ve^ and *Rbfox3*^-ve^ nuclei (n=15 nuclei from 2-3 animals/group). \*\*\*\**P* < 0.0001. **(C)** Mean fluorescence intensity (arbitrary units) of *Dory* in hippocampal subregions and cerebellum (n=2-3/group).

### Brain-specific annotation of *Dory* transcripts by high resolution RNA sequencing

The transcript structures of many, if not most, lncRNAs are not well characterized because of insufficient depth of sequencing of tissues expressing region- or cell-specific lncRNAs [58]. As of August 2024, when this study was initiated, the NCBI database displayed a single two-exon transcript for *Dory* (*2700046G09Rik*/*Sgms1os1*) on chromosome 19, whilst Ensembl listed 4 transcripts of 1-3 exons. We performed RNA capture sequencing to establish *Dory* transcript models in mouse brain (Fig. 3). Capture sequencing samples contained normalized *Dory* gene counts (count / total reads × 1,000,000) (Fig. 3A) of ∼70-150 compared to ∼2-3 counts in non-capture samples, representing an average enrichment of ∼40-fold (Fig. 3B). In addition to the 18 *Dory* transcript isoforms now listed in Ensembl, our capture sequencing identified a novel variant (Fig. 3C), similar in structure to Ensembl variant ENSMUST00000181612 (*Dory* V2), with two exons but starting from the same start site as *Dory* V4, with an extended first exon. Comparison to the human database identified a possible human ortholog, *SGMS1-AS1*, located on chromosome 10 with 4 exons (Fig. 3D). Expression of all transcripts was not significantly different between sexes (Fig. 3E).

**Fig. 3:**
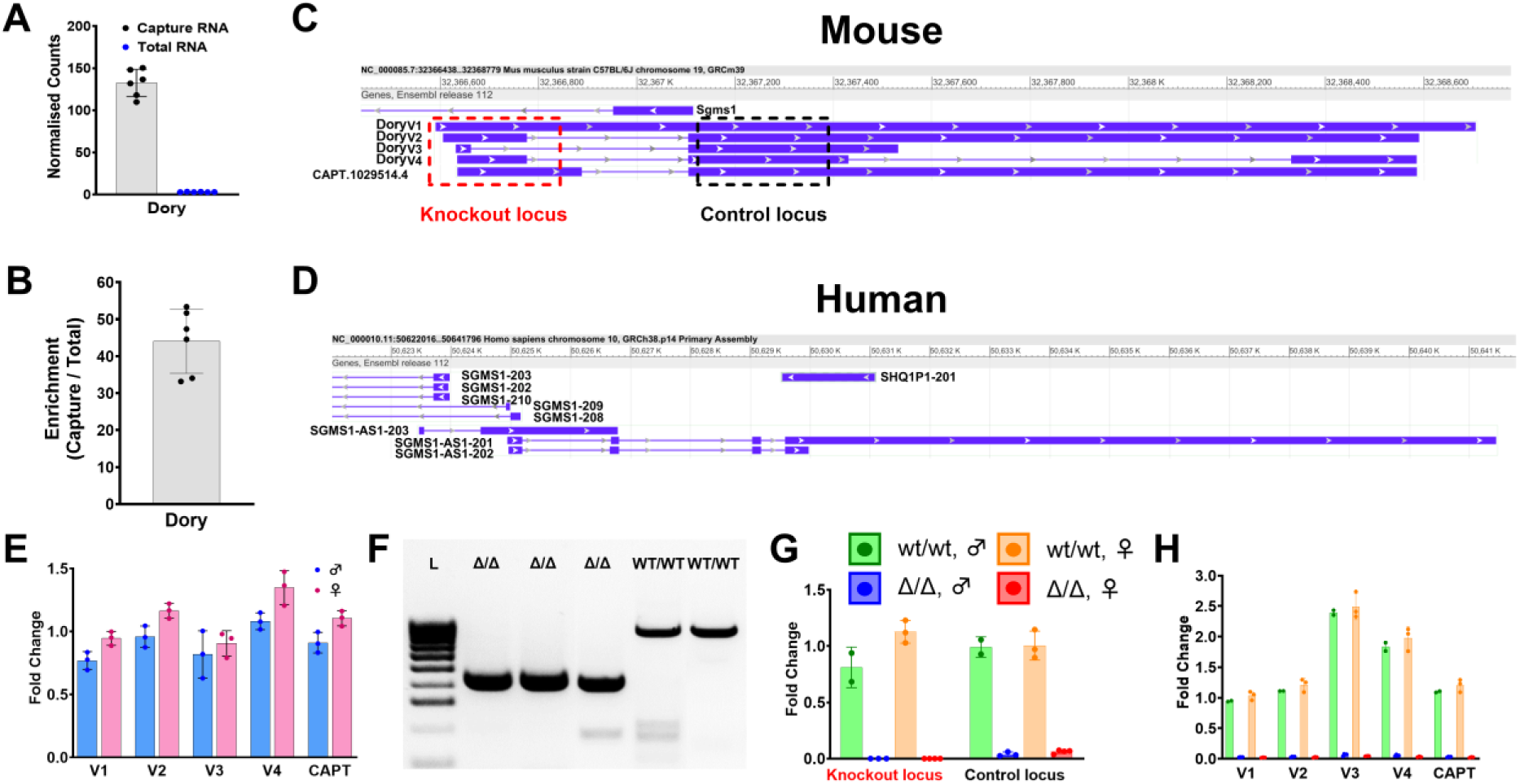
RNA capture sequencing and knockout of *Dory* (*270046G09Rik*/*Sgms1os1*). **(A)** Normalized RNA sequencing gene counts for capture sequencing and total uncaptured brain RNA samples (n=6/group). **(B)** Enrichment of capture samples (n=6). **(C)** UCSC genome browser track of the *Dory* gene locus showing the transcript models for annotated transcript variants and the novel variant found by capture sequencing (CAPT.1029514.4). The red box indicates the locus removed by CRISPR-Cas9 gene editing. The black box indicates the locus selected for comparison in **(G)**. **(D)** UCSC track of *SGMS1-AS1*, the possible human ortholog of *Dory*. **(E)** Expression of *Dory* transcript variants in RNA capture sequencing samples (n=3/group). Fold change is relative to average expression of all variants. **(F)** Representative genotyping by gel electrophoresis of wild type (wt/wt) and knockout (Δ/Δ) DNA. **(G)** *Dory* transcript expression of the knockout locus and control locus in knockout (Δ/Δ) and wild type (wt/wt) hippocampus by quantitative PCR (n=2-4/group). Fold change is relative to the average wild type expression for each locus. **(H)** Expression of *Dory* transcript variants in knockout and wild-type hippocampus (n=2-4/group). Fold change is relative to average expression of all variants. Legend in **(G)** holds for **(H)**.

### Generation of *Dory* null mice by CRISPR-Cas9 deletion

To study *Dory* function *in vivo*, we generated CRISPR-Cas9-mediated knockout (Δ/Δ) mice, using guide RNAs designed to target exon 1, which is constitutive (Fig. 3C). Gene locus amplification by PCR (Fig. 3F) and Sanger sequencing (Supplementary Fig. S3) confirmed successful knockout of the exon. Quantitative PCR using primers for exon 1 or exon 2 confirmed absence of *Dory* transcripts in knockout mice (Fig. 3G). More detailed expression analysis confirmed absence of all *Dory* transcript variants in hippocampal RNA samples from knockout mice (Fig. 3H).

### Brain-specific annotation of *Dory* transcripts by high resolution RNA sequencing

The transcript structures of many, if not most, lncRNAs are not well characterized because of insufficient depth of sequencing of tissues expressing region- or cell-specific lncRNAs [58]. As of August 2024, when this study was initiated, the NCBI database displayed a single two-exon transcript for *Dory* (*2700046G09Rik*/*Sgms1os1*) on chromosome 19, whilst Ensembl listed 4 transcripts of 1-3 exons. We performed RNA capture sequencing to establish *Dory* transcript models in mouse brain (Fig. 3). Capture sequencing samples contained normalized *Dory* gene counts (count / total reads × 1,000,000) (Fig. 3A) of ∼70-150 compared to ∼2-3 counts in non- capture samples, representing an average enrichment of ∼40-fold (Fig. 3B). In addition to the 18 *Dory* transcript isoforms now listed in Ensembl, our capture sequencing identified a novel variant (Fig. 3C), similar in structure to Ensembl variant ENSMUST00000181612 (*Dory* V2), with two exons but starting from the same start site as *Dory* V4, with an extended first exon. Comparison to the human database identified a possible human ortholog, *SGMS1-AS1*, located on chromosome 10 with 4 exons (Fig. 3D). Expression of all transcripts was not significantly different between sexes (Fig. 3E).

### Generation of *Dory* null mice by CRISPR-Cas9 deletion

To study *Dory* function *in vivo*, we generated CRISPR-Cas9-mediated knockout (Δ/Δ) mice, using guide RNAs designed to target exon 1, which is constitutive (Fig. 3C). Gene locus amplification by PCR (Fig. 3F) and Sanger sequencing (Supplementary Fig. S3) confirmed successful knockout of the exon. Quantitative PCR using primers for exon 1 or exon 2 confirmed absence of *Dory* transcripts in knockout mice (Fig. 3G). More detailed expression analysis confirmed absence of all *Dory* transcript variants in hippocampal RNA samples from knockout mice (Fig. 3H).

### Deletion of *Dory* in mice impairs spatial memory formation

Male and female knockout (Δ/Δ) mice did not exhibit any overt physical or developmental phenotypes, and their hippocampi were morphologically indistinguishable from wild type (Fig. 4A). As a major function of the hippocampus is memory formation and maintenance [59], we assessed whether lack of *Dory* expression alters hippocampal memory, using spontaneous exploration behavior testing (open field, OF) followed by probing spatial (object location test, OLT) and non-spatial (novel object recognition test, NOR) memory (Fig. 4B).

**Fig. 4:**
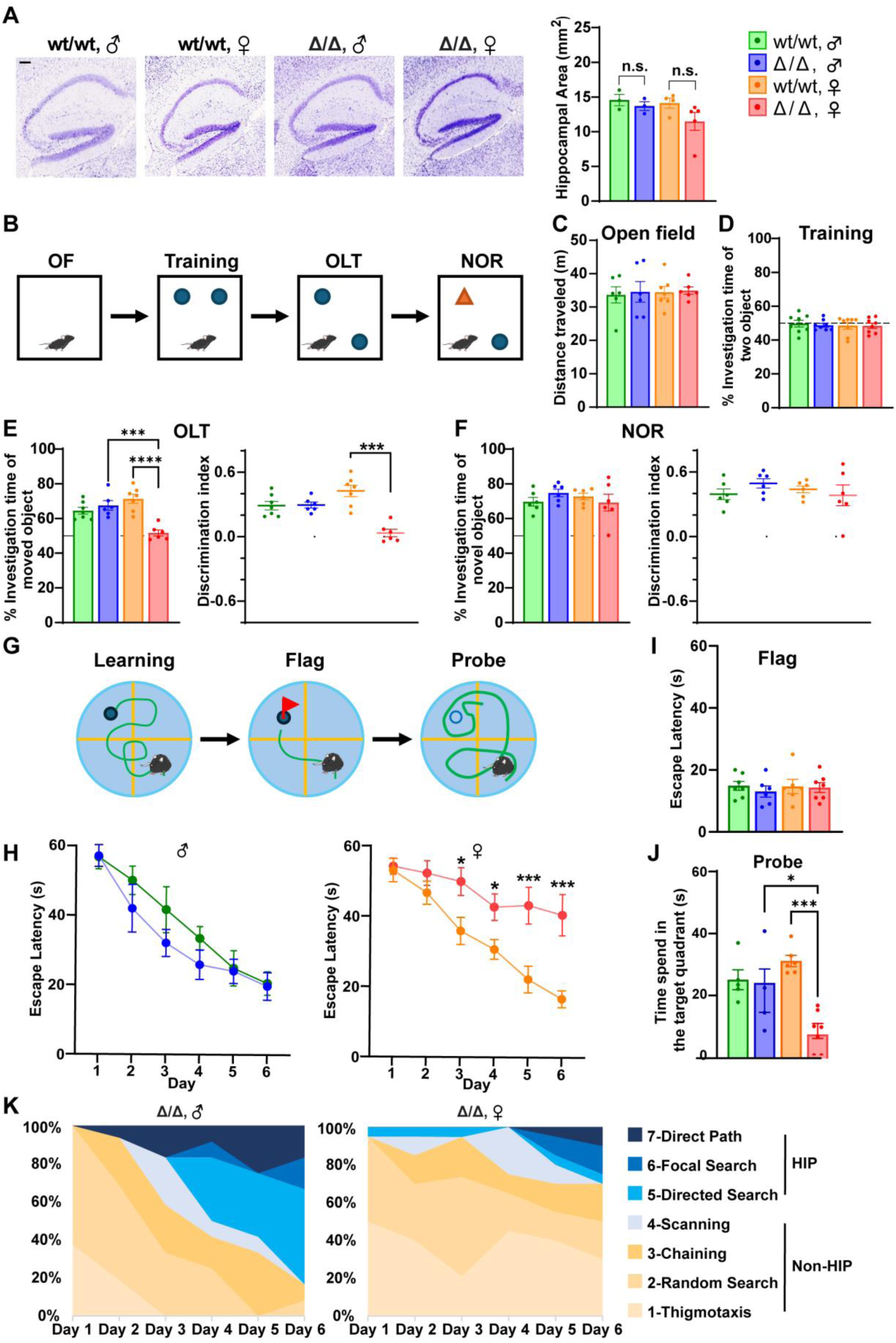
Deletion of *Dory* (*2700046G09Rik*/*Sgms1os1*) in mice significantly impairs spatial memory. **(A)** Representative images and quantitative analysis of Nissl staining of hippocampi in *Dory*^wt/wt^ and *Dory*^Δ/Δ^ mice (n=3-4/group). Scale bar = 200 µm. **(B)** Schematic representation of locomotor activity (open field test, OF), spatial memory (object location test, OLT) and non-spatial memory (novel object recognition test, NOR). **(C)** No significant difference in locomotor activity in open field test (n=6/group). **(D)** No significant difference in object investigation time during OLT and NOR training (n=8/group). **(E)** Results for moved object investigation during OLT trials (n=6-7/group) and **(F)** novel object investigation during NOR trials (n=6-7/group) by analyzing investigation time (left) and discrimination index (right). **(G)** Schematic representation of spatial memory and learning using the Morris water maze test. **(H)** Significantly longer escape latency in female Δ/Δ mice, but not in male Δ/Δ mice, reveals impaired spatial learning in a sex-dependent manner (n=5-10/group). **(I)** Escape latencies were not different in sight of flag (n=6-7/group). **(J)** Memory formation assessed with probe trials after removing the platform shows significantly lower time spent in the target quadrant for female Δ/Δ but not in male Δ/Δ compared to wt/wt (n=4-7/group). \**P* < 0.05, \*\*\**P* < 0.001, \*\*\*\**P* < 0.0001. **(K)** Qualitative swim path analysis of spatial learning shows impaired hippocampus-dependent memory formation in female Δ/Δ mice. Legend holds for all graphs excluding **(K)**.

Both WT and Δ/Δ mice exhibited similar locomotor activity in the open field test (Fig. 4C) and showed no significant preference for objects during the training phase for subsequent memory testing (Fig. 4D). However, WT mice, regardless of sex, and male Δ/Δ mice exhibited increased exploration of the object in its new location during OLT, whereas female Δ/Δ mice showed no preference, as evidenced by a significantly lower discrimination index in Δ/Δ females (Fig. 4E). This finding suggests spatial memory deficits in female Δ/Δ mice [60]. The NOR utilizes a number of brain regions beyond the hippocampus, including cortical structures such as the entorhinal, perirhinal and parahippocampal cortices [61], and subcortical structures such as the subiculum and thalamus [62]. Both WT and Δ/Δ mice, irrespective of sex, displayed equivalent interest in novel objects, demonstrating the absence of general memory impairments (Fig. 4F).

Formation of spatial memory was then assessed in more detail using the Morris water maze (MWM) test (Fig. 4G). Here, male Δ/Δ mice demonstrated comparable spatial learning to WT, successfully locating the submerged escape platform using visual cues and recalling the target quadrant (TQ) during subsequent probe trials with the platform removed (Fig. 4J). Conversely, females exhibited significantly delayed spatial learning, which did not improve beyond day 4, and consequently poor recollection of the target location during probe trials (Fig. 4H,J). All experimental groups were able to locate the platform when it was marked by a flag (Fig. 4I), suggesting no overt visual or motor deficits. A detailed analysis of swim patterns, by categorizing patterns into hippocampus- or non-hippocampus-dependent pathways during the learning phase [63, 64], revealed that Δ/Δ males progressively adopted hippocampus-dependent search strategies, but Δ/Δ females failed to do so (Fig. 4K). We named the lncRNA *2700046G09Rik ‘Dory’* due to its sex-specific effects in reference to the well-known animated movie character with memory deficits.

Interpreting genomic sequence deletions is difficult as mammalian genes are not discrete, but rather present as a continuum of overlapping and interlacing transcripts from both strands [65–68]. While the targeted sequences avoided *Sgms1* exons, it resulted in partial removal of the *Sgms1* and *Dory* (*2700046G09Rik*/*Sgms1os1*) promoters (Supplementary Fig. S4A-C). ReMap CHIP-seq data indicate two transcriptional regulator binding peaks within this region, one around the start site of annotated *Sgms1* transcripts and another around the start site of *Dory* transcripts, both within the *Sgms1* promoter region annotated by Ensembl (Supplementary Fig. S4B, C). The deletion only covers the peak associated with the *Dory* transcriptional start site (Supplementary Fig. S4B). *Sgms1* encodes an enzyme (Sphingomyelin Synthase 1) in the sphingosine-1-phosphate (S1P) signaling pathway, which regulates sphingomyelin synthesis, and which Allen Brain Cell Atlas data shows is ubiquitously expressed in the hippocampus whereas *Dory* expression is enriched in neurons (Supplementary Fig. S5). While there was a near complete loss of *Dory* expression (Fig. 3), there was only a mild (2-fold) reduction in *Sgms1* RNA expression in the knockout mice (Supplementary Fig. S4D). As genes encoding enzymes are rarely haploinsufficient [69], there is no evidence of phenotypes in *Sgms1* heterozygous mice [70], and none of the phenotypes associated with lack of *Sgms1* (tremors, infertility, decreased body weight (Supplementary Fig. S6) and premature death [56]) were observed in *Dory* knockout mice, it is unlikely that the reduction in *Sgms1* expression is responsible for the cognitive and motor phenotypes that we observed.

### ASO knockdown of *Dory* in rats impairs spatial memory

To confirm knockout spatial memory phenotypes were due to perturbance of *Dory* RNA and not DNA elements, we performed ASO-mediated knockdown of *Dory* by direct injection into male and female (8/group) rat dorsal hippocampus. Preliminary experiments suggested a mix of ASOs was most effective and that knockdown of *Dory* could be detected 1 week after injection (Supplementary Figure S7A). Knockdown was detected in rats used for behavioral tests ∼4 weeks post-injection (Fig. 5A). Like mouse knockouts, we did not detect differences in the OF (Fig. 5B and Supplementary Fig. S7B). We found support for mouse OLT results with differences in the discrimination index in females but not males (Fig. 5C). While female controls showed a preference for exploring the object in the novel location, knockdown females did not. Rat knockdown NOR results supported mouse knockouts, with no difference in discrimination index (Fig. 5D), although there was greater total exploration in female knockdown during training (Supplementary Fig. S7C). We could not replicate the mouse MWM results with the Barnes maze in knockdown rats (Supplementary Fig. S7D, slight improvement in search strategy score in knockdown rats). This may be due to differences between species (greater variability in outbred rat strain compared to inbred mouse strain), developmental effects (genetic knockdown compared to knockdown in adults), and/or technical effects of partial knockdown compared to knockout.

**Fig. 5:**
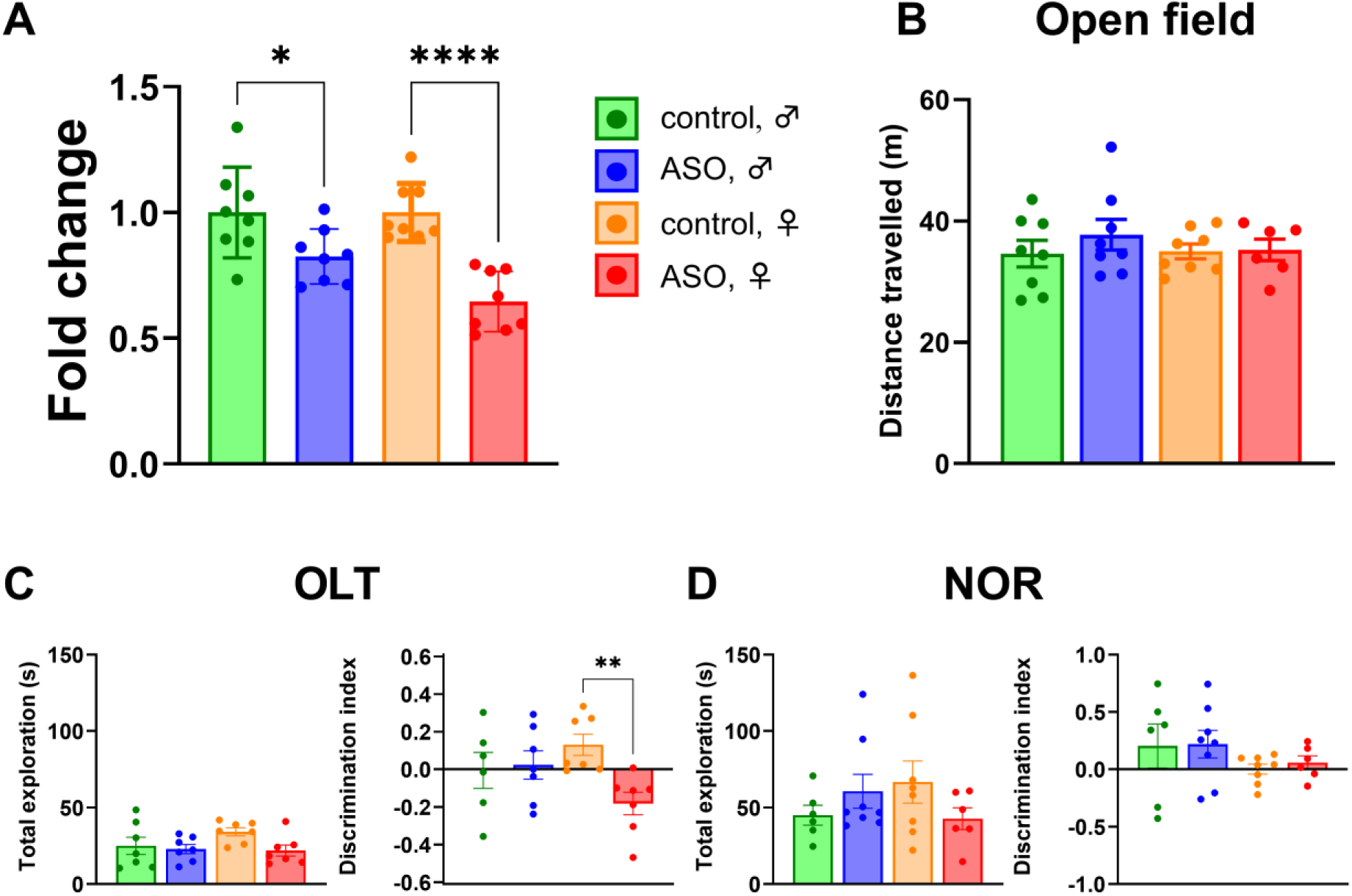
Knockdown of *Dory* in rat dorsal hippocampus significantly impairs spatial memory. **(A)** Knockdown of *Dory* in dorsal hippocampus confirmed by qPCR. **(B)** Open field test total distance travelled in knockdown rats. **(C)** Object location test (OLT) and **(D)** novel object recognition test (NOR) total exploration (left) and discrimination index (right). Female controls showed a preference for the novel location in the OLT, but female knockdowns did not. \**P* < 0.05, \*\**P* < 0.01, \*\*\*\**P* < 0.0001. Legend holds for all graphs.

### Deletion of *Dory* in mice impairs balance

The cerebellum of *Dory* Δ/Δ mice showed no morphological difference in both male and female compared to the cerebellum of WT mice (Fig. 6A). To evaluate *Dory*’s cerebellar functions, we implemented a comprehensive battery of motor activity tests. Initially, assessments in the OF and MWM tests revealed no overt motor impairments (Fig. 4C, I). Similarly, grip testing was unaffected suggesting normal muscle strength (Fig. 6B). The pole test, demanding more intricate motor skills, unveiled a subgroup of females struggling with descent (n=4 of 10), although their completion times were comparable when successful (Fig. 6C). Although Δ/Δ mice showed slightly slower descent times, the difference was marginal (Fig. 6D). To gain a more detailed perspective, we employed the rotarod test for three consecutive days (Fig. 6E, F). The latency to fall from the rotating rod was shorter in female Δ/Δ compared to male Δ/Δ and both sexes of wild-type mice, indicating declined motor control in female Δ/Δ mice. Finally, the beam test, evaluating motor coordination, exposed balance deficits exclusive to female Δ/Δ mice (Fig. 6, H). In conclusion, while the knockout of *Dory* did not appear to impact muscle strength, it evidently compromises more complex motor tasks involving balance and coordination, virtually exclusively in females. The effects on spatial memory, wholly restricted to females, further highlight the sex-specific nature of *Dory*.

**Fig. 6:**
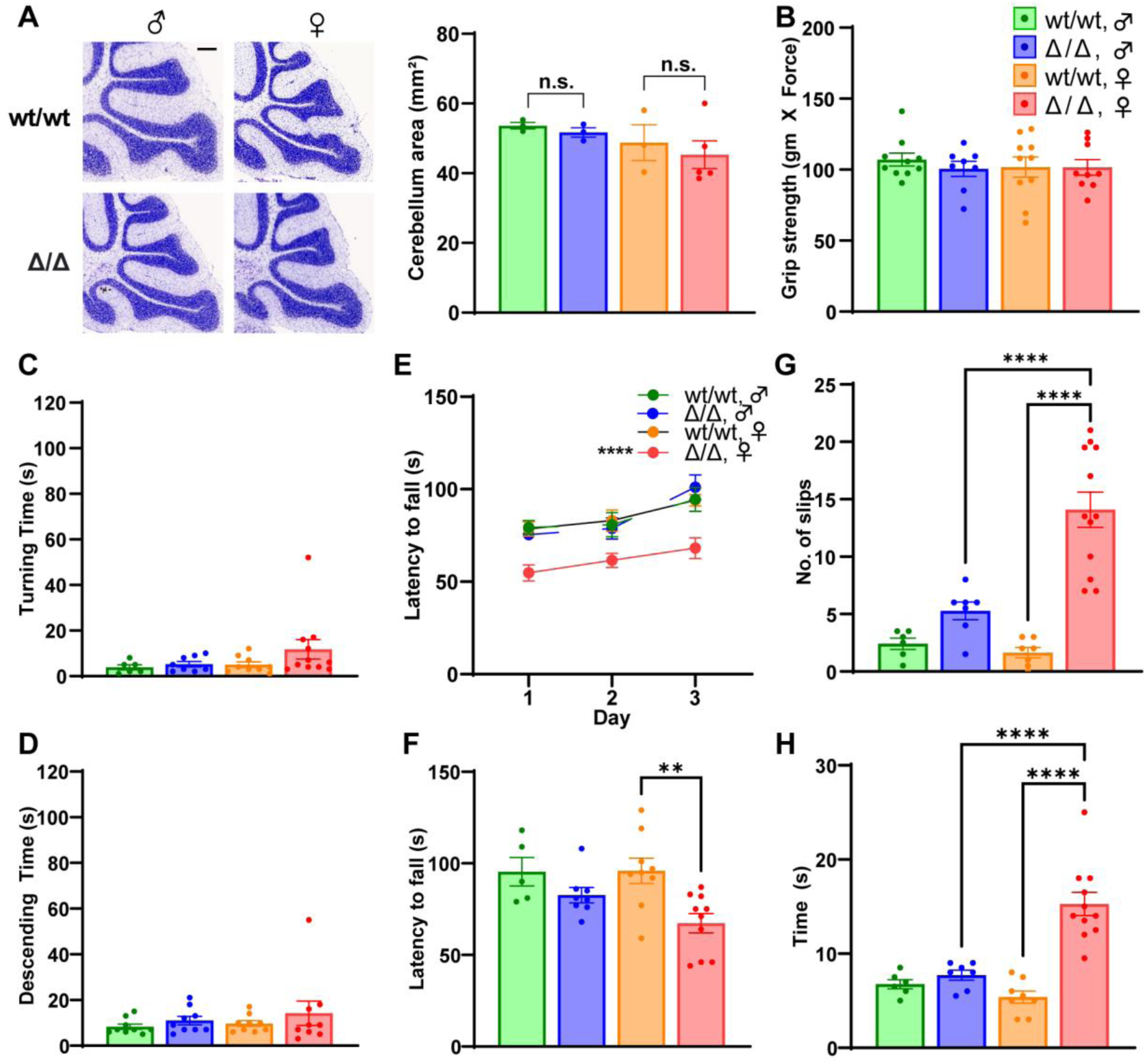
Impairment of motor balance after *Dory* deletion in mice. **(A)** Representative images and quantitative analysis of Nissl staining of cerebellum in *Dory*^wt/wt^ and *Dory*^Δ/Δ^ mice (n = 3- 5/group). Scale bar = 200 μm**. (B-H)** Measurement of motor control after deletion of *Dory* in 3-month-old mice using **(B)** grip test (n=8-10/group), **(C,D)** pole test (n=6-11/group, **(E,F)** rotarod test (n=5-10/group) and **(G,H)** beam test (n=6-11/group). Intact motor strength and locomotor activity were observed in the **(B)** grip and **(C,D)** pole tests. Deletion of *Dory* impairs female motor control and balance in the **(E,F)** rotarod and **(G,H)** beam tests. **B-D** and **G-H** measurements were made on day 3. \*\**P* < 0.01, \*\*\*\**P* < 0.0001. Legend holds for all graphs.

### Dory loss induces sex-dependent transcriptional and proteomic changes in the hippocampus

To characterize the molecular changes associated with the knockout of *Dory*, we performed RNA sequencing to measure transcriptome-wide gene expression, and mass spectrometry to assess proteome-wide changes in protein expression (Supplementary Fig. S8). Differential expression analysis (all ≥ 2-fold, p-value < 0.05) in female Δ/Δ mice identified 70 genes, of which 14 were downregulated and 56 upregulated, and 74 proteins, of which 49 were downregulated and 25 upregulated (Fig. 7A). Differential expression analysis in male Δ/Δ mice identified 684 genes, of which 224 were downregulated and 462 upregulated, and 107 proteins, of which 73 were downregulated and 34 upregulated (Fig. 7A). RNA sequencing and mass spectrometry results are available in Supplementary Files S1-S3.

**Fig. 7:**
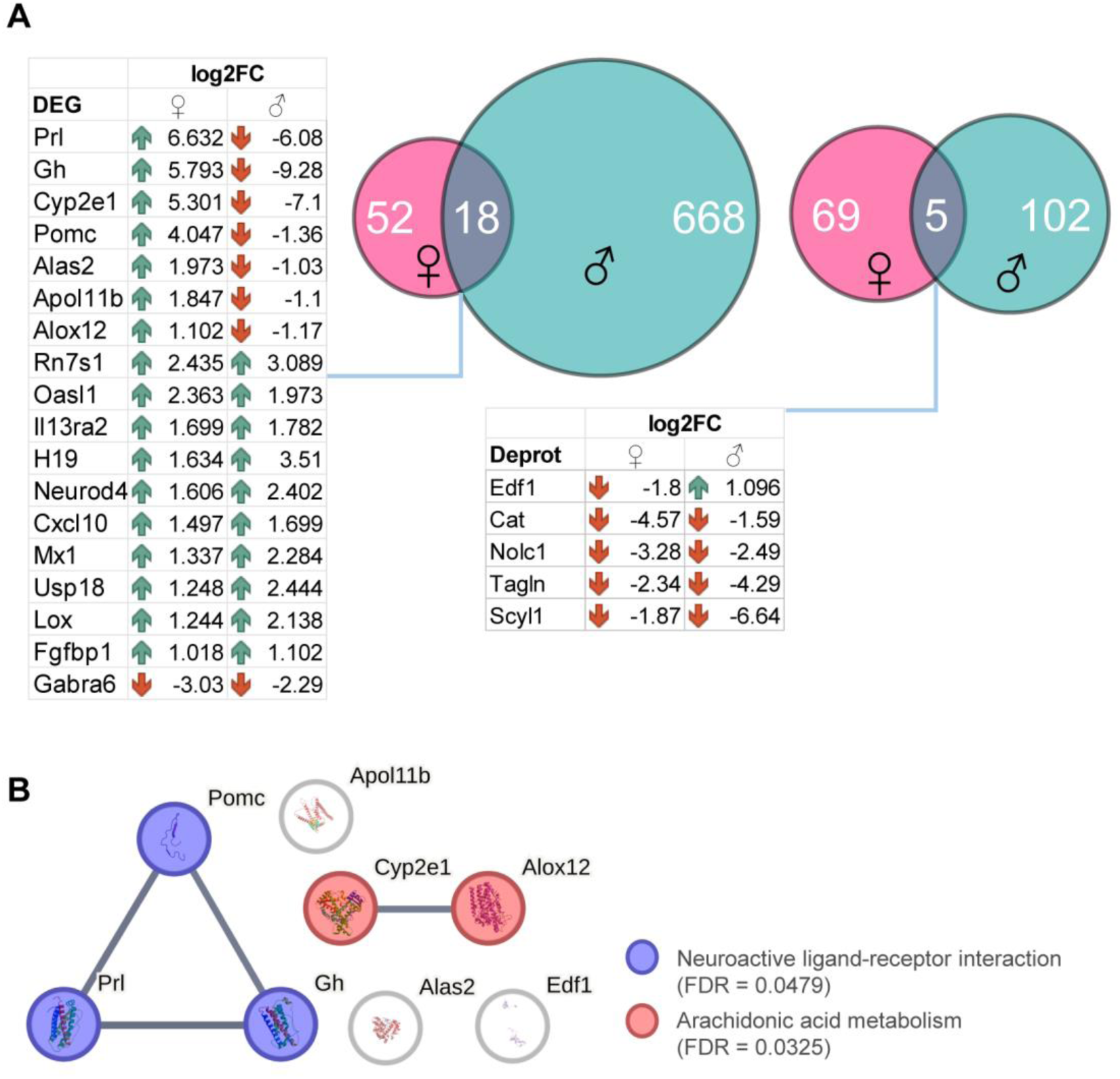
Sex-dependent molecular responses to Dory loss in the hippocampus. **(A)** List and Venn diagrams of overlapping differentially expressed genes (left) and proteins (right) and their overlap between male and female mice after *Dory* knockout. **(B)** Protein-protein interaction network of 8 differentially expressed genes, analyzed using the Search Tool for the Retrieval of Interacting Genes/Proteins database (STRING). Enrichment analysis by KEGG pathways is shown (FDR < 0.05).

Eighteen genes and five proteins were found to be differentially expressed in both male and female Δ/Δ hippocampi in the RNA-seq data, compared to WT mice (Fig. 7A). Most genes were upregulated in female Δ/Δ mice, while 7 genes (*Prl, Gh, Cyp2e1, Pomc, Alas2, Apol11b* and *Alox12*) were upregulated in female Δ/Δ mice but downregulated in male Δ/Δ mice. In mass spectrometry data, Edf1 was the only differentially expressed protein in male (up) and female (down) Δ/Δ mice hippocampi with opposing directionality. Vps30 and Kcnq4 were markedly increased in female Δ/Δ hippocampi, while Cbl and Gc were markedly decreased. Protein-protein interaction analysis of commonly dysregulated genes using STRING revealed that Prl, Gh and Pomc are associated in females via a neuroactive ligand-receptor interaction likely involved in neurogenesis [71], while Cyp2e1 and Alox12 have a shared association with arachidonic acid metabolism (Fig. 7B).

We then performed a functional enrichment analysis of differentially expressed genes and proteins in the hippocampi of female and male *Dory* Δ/Δ mice (Supplementary Fig. S8). The biological processes enriched among differentially expressed genes in female Δ/Δ mice were generally associated with cellular responses to cytokine stimulus and hormone secretion (Supplementary Fig. S8B), while differentially expressed genes in male Δ/Δ mice identified the mitotic cell cycle (Supplementary Fig. S8C). Interestingly, a subset of female Δ/Δ differentially expressed genes including *Gh* are associated with cognitive deficits [72].

## Discussion

Our behavioral data shows that deletion of *Dory* results in female-specific deficits in memory formation and motor coordination. *Dory* deletion reduced hippocampus-dependent spatial learning and spatial memory formation in female mice, but did not compromise non-spatial memory formation in either sex. Rat knockdown experiments support that *Dory* (not DNA elements in the locus) is responsible for the spatial memory deficits. Our findings identify *Dory* as a regulator of sex-specific hippocampus-dependent cognitive functions.

The hippocampus is essential not only for memory but also for the spatial representation of environments and navigation [59]. In rodents, a number of sex differences in spatial memory and navigation task performance have been noted (see [73]). In the MWM, male rodents are typically faster in learning the platform location in training [74–77] and use more spatial strategies during the first training days. Importantly, male and female rodents typically navigate using different spatial navigation strategies, with males preferring place strategy and females preferring response/cue strategy [78, 79]. Strategy choice appears to be related to cholinergic muscarinic receptor binding in the (predominantly ventral) hippocampus [80], and the choice of strategy has classically been associated with hippocampal (place) or striatal (response/cue) activation [78]. Females also seem to be more sensitive to cholinergic blockade when performing spatial tasks [81], while males have larger long-term potentiation at perforant pathway-dentate gyrus (PP-DG) synapses [74].

There are well-documented differences in spatial learning and memory between males and females in humans, where (on average, with high variance) males exhibit stronger spatial navigation and orientation abilities [74, 82, 83]. Sex differences in the virtual MWM task have been associated with hippocampal/parahippocampal theta wave power during navigation of a familiar environment, and high-gamma power during inter-trial rest periods after navigating a new environment [84]. Together, these results suggest fundamental differences in the cognitive programming of spatial navigation behaviors between the sexes. While all hippocampal subregions had intact morphology in both sexes, we cannot exclude that there may be subtle differences as males and females have different synaptic densities in the hippocampus [85].

Our RNA sequencing data showed that the knockout of *Dory* increased expression of genes associated with hormones and hormone secretion in the hippocampus of female but not male mice, including prolactin, POMC and growth hormone, suggesting the presence of hormone and sex-specific transcriptional regulatory responses in the hippocampus. High levels of prolactin are associated with cognitive impairment [86] and associated with cognition in female patients with prolactinomas [87]. Prolactin can also modify hippocampal gene networks [88]. Growth hormone (GH) is known to enhance cognitive impairment [89], and high postnatal GH levels may contribute to neurodevelopmental abnormalities [90]. Furthermore, navigational strategies in males [91] and females [92] are influenced by sex hormone status, preferring place strategy when high and cue strategy when low. Differences in the expression and secretion of hormones between sexes suggests that aspects of the sex-specific deficits in spatial memory formation and balance may be hormonally related, especially since synaptic density in the female brain is affected by hormonal fluctuations [85].

Thirty-two differentially expressed genes in the female hippocampus were associated with cellular response to cytokines, a biological process which is associated with cognitive impairments during neurological dysfunction [93]. Many of these cytokines have been implicated in components of hippocampal memory. For example, Gipc1 associates with both surface and cytoplasmic NMDA receptors in hippocampal neurons [94], and Prrt1 is required for NMDA receptor-dependent long-term depression and proper NMDA-induced AMPAR trafficking [95], with knockout mice displaying cognitive deficits in the MWM and novel object recognition tasks [96]. Furthermore *Smpd3* deficiency perturbs neuronal proteostasis and causes progressive cognitive impairment [97], Kpnb1 is found within cytoplasmic granules in hippocampal neurons in Alzheimer’s disease and co-localizes with hyperphosphorylated tau [98], *Grin2b* is necessary for normal neuronal development and is important for learning and memory [99], and Vcam1 labels a subpopulation of neural stem cells in the hippocampus and contributes to spatial memory [100].

It has been reported that *Dory* regulates neuronal myelination by modulating the expression of the nearby *Pten* gene in a *microRNA-23a*-dependent manner [101]. The human ortholog of *Dory* (*SGMS1-AS1*) has been associated with cancer progression via a *microR-106a-5p*/MYLI9 axis [102]. However, the *Pten* gene was not differentially expressed in the hippocampus of *Dory* knockout mice of either sex.

We suggest that *Dory* contributes to negative transcriptional regulation of target genes, as most differentially expressed genes were upregulated in the absence of *Dory*. Analysis of *Dory* distribution in the hippocampus using RNAscope demonstrated that it has a punctate subnuclear localization, which suggests a role for *Dory* in transcription/enhancer function. Furthermore, the only differentially expressed protein that had opposing effects in males (up) and females (down), Edf1, is a transcriptional coactivator [103], suggesting that *Dory* may influence transcriptional networks through interactions with transcriptional coactivator complexes. Further studies are required to determine whether *Dory* directly interacts with transcriptional regulators or chromatin-associated complexes to control transcriptional programs. Mass spectrometry also revealed that the expression of Vps30 (a component of the endosome recycling system) and Kcnq4 (a potassium channel thought to regulate neuronal excitability) were increased in knockout females, whereas Cbl (an E3 ubiquitin-protein ligase) and Gc (a vitamin D-binding and transport protein) were both decreased in knockout females. Although the significance of these protein changes remains unclear, several of the affected proteins are involved in processes related to neuronal signaling, membrane trafficking and protein turnover, suggesting broader alterations in neuronal regulatory pathways following *Dory* deletion.

Examination of the functional roles of *Dory* in the cerebellum found that balance impairment occurred in female but not male knockout mice. RNAscope analysis revealed that *Dory* is not only expressed in the cerebellum granular layer neurons, but also in other *Rbfox3*^+ve^ neuronal cells. Biological sex-related differences in body coordination by hormonal factors and neuromuscular control have been reported [104], but the underlying molecular mechanisms are not understood.

Two other lncRNAs have been reported to be required for spatial learning, examined in mixed gender cohorts with no observed sex differences [41], or only in male mice [42]. On the other hand, two lncRNAs have been shown to have differential expression and/or affect non-memory behavior differently in males and females. The human lncRNA *FEDORA* is upregulated in the prefrontal cortex of females but not males suffering severe depression, and its ectopic expression in mice resulted in depression-like behavior in females only [49, 50]. In male mice, knockout of *Pnky*, a neural lncRNA that regulates brain development, resulted in deficits in cued fear recall, whereas in female mice the acoustic startle response (ASR) was increased and accompanied by a decrease in prepulse inhibition, both of which are behaviors altered in affective disorders. Ectopic expression of *Pnky* reversed the ASR phenotype of female *Pnky* knockout mice [105]. Together with our findings of female-specific spatial memory regulation by *Dory*, a picture is emerging of lncRNAs regulating specific neuronal functions in a sex-specific manner.

There are many other lncRNAs that show differential expression between males and females [106–108], although most have not been functionally characterized, and regrettably many studies of effects of lncRNAs on cognitive functions have been conducted only in male mice (e.g., [30, 109–111]). Our findings highlight the importance of considering biological sex when investigating the roles of brain-expressed lncRNAs. More broadly, they suggest that lncRNAs may act as regulators of sex-specific transcriptional programs that shape neural circuit function and behavior.

## Materials and Methods

### Animals

Healthy 3-month-old C57BL/6 (WT and/or Δ/Δ) male and female mice were used for all experiments except ASO knockdown. Mice were maintained under standard housing conditions, on a 12-hour light-dark cycle, with food and water available *ad libitum*. All mice were euthanized by cervical dislocation and transcardially perfused with 50 mL phosphate buffered saline (1X PBS) before tissue collection.

Healthy 2.5-month-old Long-Evans male and female rats were used for ASO experiments, with females weighing 220-260 g and males weighing 280-420 g. Rats were obtained and bred in-house at the University of New South Wales (Kensington, NSW, Australia) and housed in groups of four in opaque plastic cages (1500U, Tecniplast) with a floor area of 1500 cm² (L 51.3 × W 38.1 × H 25.6 cm). The boxes were kept in an air-conditioned colony room maintained on a 12-hour light-dark cycle (lights on at 7:00 p.m.). Food and water were available *ad libitum* in the home cage during all phases of the experiment. All experimental procedures occurred between 8:00 a.m. and 6:00 p.m. All rats were euthanized via an intraperitoneal injection of pentobarbitone solution (Lethabarb; 325 mg/mL per rat).

All mouse work was undertaken with the approval of the Macquarie University Animal Ethics Committee (Ethics numbers ARA 2017/033, ARA 2018/019, ARA 2018/020). Rat experiments were undertaken with the approval of the University of New South Wales Animal Ethics Committee (Ethics number iRECS7890). All animal experiments were conducted in accordance with New South Wales and Australian animal research guidelines.

### Generation of knockout mice and genotyping

*Dory* (*2700046G09Rik)* knockout mice were generated on the C57BL/6J background by CRISPR-Cas9 gene editing. Four single guide RNAs (sgRNAs) (5′-TGCCTGGGGCTGCGCGATTT-3′, 5′-TTCCTAAATCGCGCAGCCCC-3′, 5′-TCCGGGGCCGTAGACGAAGA-3′, 5′-CCCGGACCCTTCTTCGTCTA-3′), targeting the first exon of *Dory* (*ENSMUSG00000097787*) were rationally designed using a computational tool to minimize potential off-target effects [112] and incubated with a high fidelity Cas9 protein (*Alt-R* HiFi Cas9 Nuclease, IDT) to form ribonucleoprotein (RNP) complexes. RNPs were delivered to fertilized zygotes by electroporation (Nepa21, Nepagene). The embryos were then implanted to foster females following conventional methodology [113]. Mice were genotyped by amplicon PCR and agarose gel electrophoresis. Primers used for mouse genotyping by PCR were 5′-GCAAAAGGCGGCTAGAAGGT-3′ and 5′-TTAGCCGTTAGGTTCTGGCTC-3′.

Two founder F0 mice were selected and intercrossed to generate F1 (*Dory*^−/−^) mice for breeding of knockouts. Homozygous (*Dory*^−/−^) mice were crossed with C57BL/6J wild-type mice to generate heterozygous (*Dory*^+/−^) mice, and control wild-type (*Dory*^+/+^) mice were obtained by intercrossing heterozygous (*Dory*^+/−^) mice. The targeted deletion of ∼430 bp encompassing exon 1 (containing the transcriptional start) of the *Dory* locus was confirmed by Sanger sequencing of amplicons of target DNA generated by PCR using genotyping primers in F2 mice (Supplementary Fig. S1). Sequences generated by Sanger sequencing were aligned to the GRCm39 mouse genome using NCBI blastn with default settings [114].

### *In situ* hybridization

Whole mouse brains were fixed in 4% PFA overnight and embedded in OCT compound and cut into 8 μm thick sagittal sections. Frozen sections were thaw-mounted onto the slides. The sections were fixed in 4% PFA for 10 min, dehydrated in 50%, 75%, 95%, and 100% ethanol for 5 min each, and finally air-dried. Tissues were then treated with pre-treatment solution for 10 min at room temperature. For RNA detection, samples were incubated with *Dory* and *Rbfox3* probes for 2 h at 40 °C, and then different amplifier solutions from the RNAscope Multiplex Fluorescent Reagent Kit (3203100) were added sequentially following the manufacturer’s instructions. The signals were coupled with Opal520 for *Rbfox3* and Opal570 for *Dory*. The RNAscope assay reagents, *Rbfox3* (313311-C3) mRNA target probe and a custom designed *Dory* (*2700046G09Rik*, 561611-C1) lncRNA target probe were purchased from Advanced Cell Diagnostics, Bio-Techne. 4′-6-Diamidino-2-phenylindole (DAPI) was used for nuclear staining. The slides were washed, mounted, and imaged on a slide scanner AxioScan (Zeiss) or a confocal laser scanning microscope (LSM880; Zeiss).

For RNAscope quantification, 3 fields of view were randomly selected in each region. For *Rbfox3*^+ve^ and *Rbfox3*^-ve^ nuclei quantification, *Dory* expression (number of dots) was quantified in equal numbers of *Rbfox3*^+ve^ and *Rbfox3*^-ve^ nuclei. For average *Dory* expression in each CNS region, mean fluorescence intensity was quantified.

For neuronal subtype co-localization, brain sections (n=2 mice, 1 section each) were labelled with DAPI, *Dory* and neuronal subtype markers. Mice were either 1 male and 1 female (*Grin1* and *Gabrg1*), or 2 females (*Chat*, *Dcx*, *Nes*, *Pcp2*). Target probes for neuronal subtypes were *Grin1* (431611-C3, excitatory), *Gabrg1* (501401-C2, inhibitory), *Chat* (408731-C2, cholinergic), *Dcx* (478671, immature), *Nes* (313161-C3, neural stem cells), *Pcp2* (509991-C2, Purkinje cells).

### Nissl staining

Nissl staining was conducted to examine hippocampus and cerebellum morphology. Three micrometer sections mounted on superfrost PLUS slides (Epredia, J1800AMNZ) were dewaxed with xylene and rehydrated with descending series of ethanol for 5 min. Sections were then immersed in running tap water for 2 min before placing in dH_2_O for 5 min. Prior to staining, 0.1% Cresyl violet (Sigma, C5042) solution was prepared, filtered, and ripened for at least 48 h, and stored in the dark at room temperature. Sections were then treated with pre-warmed 0.1% Cresyl violet solution for 10 min at 37 °C. Slides were quickly dipped into dH_2_O before differentiating in 96% ethanol containing a few drops of glacial acetic acid for 5 seconds. Slides were placed in dH_2_O to stop the differentiation process and checked under the microscope. Differentiation steps were repeated until only the Nissl bodies and nuclei were stained. Slides were quickly rinsed in 96% ethanol before being dehydrated in 100% ethanol and xylene for 10 min. Slides were coverslipped using DPX mounting medium (Sigma-Aldrich Inc., USA, 06522) and imaged with Zeiss Axio scan Z1.

### RNA isolation and qPCR

For mouse studies, three-month-old mice were euthanized by cervical dislocation and the brains extracted. The hippocampus (HIP) and cerebellum (CB) were dissected and snap frozen in liquid nitrogen, followed by homogenization using a mortar and pestle. RNA was extracted using the RNeasy MiniKit (Qiagen, #74,106) with on-column DNase treatment (Qiagen, #79,254). Reverse transcription was performed with the SuperScript VILO cDNA synthesis kit (Invitrogen; #11754050). Quantitative real-time PCR (qPCR) was performed on QuantStudio™ 7 Pro and analyzed using Design & Analysis Software 2.7.0 (ThermoFisher, USA).

For rat knockdown experiments rats were euthanized four weeks (or 1 week for preliminary knockdown tests) after ASO injection via an intraperitoneal injection of pentobarbitone solution (Lethabarb; 325 mg/mL per rat). Immediately following euthanasia, brains were extracted and rapidly frozen in liquid nitrogen, then stored at −80 °C. For regional analysis, 1 mm thick coronal brain sections (including the injection sites) were obtained using a rat brain matrix. Micropunches of the hippocampus at the injection sites were then collected from this coronal section using a 1.0 mm disposable micropunch, guided by coordinates from the rat brain atlas. RNA was extracted using the RNeasy Plus Mini Kit (Qiagen, 74134) following the manufacturer’s instructions. Approximately 200 ng of RNA was treated with TURBO DNase (Invitrogen), reverse transcribed using the SensiFAST cDNA Synthesis Kit (Meridian), and purified with the Zymo Clean & Concentrator-25 Kit (R1018). qPCR was performed using the SensiFAST SYBR Hi-ROX Kit (BIO-92005) on a ViiA 7 Real-Time PCR System (Applied Biosystems). *Gapdh* and *Actin* were used as housekeeping controls. Data were analyzed using the ΔΔCt method in Microsoft Excel and statistical analyses were performed in GraphPad Prism (unpaired t-tests).

### RNA capture sequencing and generation of transcript models

Whole brains from 4 male and 4 female WT mice were hemi-sectioned and snap frozen in liquid nitrogen, followed by homogenization of the right brain hemisphere (3/sex) or hippocampus and cerebellum (1/sex) in lysis buffer using a motorized pestle. RNA was extracted using the RNeasy Plus MiniKit with on-column DNase treatment. RNA integrity numbers were > 9 for all samples. RNA was sent to the Ramaciotti Institute (UNSW Sydney) for library preparation, capture enrichment, and sequencing. In brief, two cDNA libraries per sample were prepared using the Illumina Stranded Total RNA Library Prep Kit according to the manufacturer’s instructions, with ribosomal RNA depletion. Following library generation, one library per sample was subjected to capture enrichment using the Twist Target Enrichment Standard Hybridization Protocol v2. Libraries were sequenced on two separate NovaSeq 6000 lanes, generating ∼47-83 million stranded 100 bp read pairs. RNA sequencing data was analysed on the high performance computing infrastructure Katana at UNSW (https://research.unsw.edu.au/katana). Data quality was checked using FASTQC [115]. Adaptors were removed using Cutadapt [116]. Forward and reverse reads were aligned using the STAR aligner [117] to the mouse GRCm39 genome from Ensembl. Total mapped reads for capture and non-capture samples were ∼484 and ∼462 million read pairs respectively. Transcriptome annotation files (GTFs) were generated for each sample using StringTie [118] with the GRCm39 Ensembl reference GFF3, and capture or non-capture GTFs were merged together.

For ASO design, we generated rat transcript models for *Dory* from a published rat brain RNA sequencing dataset [119] in the same manner as mouse models, but using the rat BN7 genome.

### RNA sequencing of hippocampi

RNA integrity numbers were > 9 for all samples. RNA was sent to the Ramaciotti Centre for Genomics (UNSW Sydney) for library preparation and sequencing. In brief, cDNA libraries were prepared using the Illumina Stranded Total RNA library preparation kit according to the manufacturer’s instructions, with ribosomal RNA depletion. Following library generation, libraries were sequenced on a single lane of a NovaSeq 6000, generating ∼108-146 million stranded 100 bp read pairs. Data quality was checked using FASTQC [115]. Adaptors were removed using Cutadapt [116]. Forward and reverse reads files were aligned using the STAR aligner [117] to the mouse GRCm39 genome from Ensembl. Total mapped reads were ∼103-139 million read pairs per sample. Gene counts were generated from BAM files using HTSeq-count [120] and assessed for differential expression between groups using Deseq2 [121]. A threshold of ±2-fold and a p-value cut-off of < 0.05 for significance were applied. Gene Ontology (GO) enrichment of differentially expressed genes was assessed using PANTHER v18 [122] or Metascape [123].

### Label free mass spectrometry of hippocampi

Hippocampal total protein extracts were prepared by sonicating ∼100 mg of fresh mouse hippocampal tissue in 1 mL of RIPA buffer (Sigma, Australia) containing cOmplete protease inhibitor cocktail (Roche, Australia). Sonication was performed on ice, using 5-6 bursts of the probe sonicator. Samples were centrifuged (∼12,000 × *g*, 6 °C, 1 h), and the supernatant transferred to clean polypropylene tubes. The total protein was assayed using the Bradford assay (BioRad, Australia), with the total protein concentration ranging between 1.5–2.5 mg/mL in each sample.

In preparation for proteomics analysis, the protein extracts (∼80 mg/40 µL) were reduced using 20 mM DTT prepared in 20 mM ammonium bicarbonate buffer (pH 8), and incubated at 37 °C for 60 min. Protein samples were then alkylated by adding 20 µL of 200 mM iodoacetamide, and incubated at ambient temperature for 15 min. A buffer exchange to 5% SDS in 100 mM ammonium bicarbonate (pH 7.5) was performed using 3 kDa spin filters (Amicon, Australia), using 3–4 washes of 250 µL of the buffer (7,000 × *g*, 1 h, 8 °C), with the final retentate volume being ∼250 µL. Next, 25 µL of 12% aqueous phosphoric acid was added to each sample (pH drops to ∼2), the solutions were transferred into 15 mL polypropylene tubes, and 3 mL of S-trap protein binding buffer was added (90% aqueous methanol containing 10 mM ammonium bicarbonate, pH 7.1). The solution was transferred to S-Trap micro spin cartridges (Protifi, Fairport NY), in as many steps as needed to pass the complete volume through the S-trap device, centrifuging at 4,000 × *g* for approximately 1 min each time. Proteins are retained on the S-trap. The S-trap and captured proteins were washed three times by adding 200 µL S-trap buffer and centrifuging at 4,000 × *g*. Captured proteins were then cleaved by tryptic digestion by transferring the S-trap to a clean tube, adding 1.5 µg of sequencing-grade trypsin (Promega, Australia) in a final volume of 230 µL of 50 mM ammonium bicarbonate, capping and sealing the S-traps with paraffin film to limit evaporative loss, and incubating at 37 °C for 16-18 h. Tryptic peptides were recovered by adding 40 µL of 50 mM ammonium bicarbonate to each S-trap followed by 0.2% aqueous formic acid (0.2%), centrifuging at each step (4,000 × *g*, ∼1 min). Hydrophobic peptides were recovered with an elution of 40 µL 50% aqueous acetonitrile containing 0.2% formic acid. All elutions were pooled, then dried using a centrifugal evaporator and resuspended in 30 µL of 1% formic acid containing 0.2% heptafluorobutyric acid (HFBA). The absorbance at 280 nm of each sample were checked using a spectrophotometer (DeNovix, Australia), and concentrations estimated using the extinction coefficient of BSA. The sample concentrations ranged between 1.4–3.7 mg/mL, indicating > 90% protein recovery for all samples.

Tryptic peptide samples were analyzed by LC-MS/MS using a high resolution accurate mass (HRAM) orbitrap mass spectrometer in ESI mode (Exploris 480 Thermo Electron, Bremen, Germany) and using data-dependent acquisition (DDA). Chromatography was carried out using a nano-LC equipped with an autosampler (Vanquish Neo UHPLC, Thermo Scientific, Waltham, USA). Peptides (∼6 µg/2 µL injected on column) were initially captured on a C18 cartridge (PepMap Neo C18, 5 mm particle size, 300 mm x 5 mm, Thermo Scientific Dionex, Waltham, USA), switching to a capillary column (25 cm) containing C18 reverse phase packing (Reprosil-Pur, 1.9 µm, 200 Å, Dr. Maisch GmbH, Ammerbuch-Entringen, Germany), supported within a column heater (45 °C, Sonation GmbH, Germany). A 120 min gradient of buffer A (H_2_O:CH_3_CN of 98:2 containing 0.1% formic acid) to buffer B (H_2_O:CH_3_CN of 20:80 containing 0.1% formic acid) was used to elute peptides, with a flow rate of 200 nL/min, with high voltage applied at the column inlet. The mass spectrometer was operated using a FAIMS device, with the following settings: ion spray voltage 2000 V, ion transfer tube temperature 280 °C, positive ion mode, data-dependent acquisition mode with a survey scan acquired (*m/z* 350-1500) and up to 20 multiply charged ions (charge state ≥ 2+) isolated for MS/MS fragmentation (intensity threshold 8.0e3). Nitrogen was used as the HCD collision gas.

Peak lists were generated from the raw files, data processed and differential proteome expression determined using ProteomeDiscoverer 2.4 software (Thermo Scientific, Waltham, USA), using a previously published approach [124]. Search parameters included: peptide precursor and MS/MS tolerances of ± 10 ppm and ± 0.05 Da respectively, dynamic modifications were acetyl N-terminus, met oxidation, phospho (ST) and phospho (Y), while carbamidomethyl cys was selected as a static modification, peptide charge of ≥ 2+, enzyme specificity trypsin with up to two missed cleavages allowed. Across-group comparisons were performed using 3-5 biological replicates, and three search engines used to maximise numbers of proteins identified (Mascot, Sequest, Amanda). For confident protein identification, a minimum of 2 unique peptides per protein were required, and an FDR of ≤ 5% and strict parsimony applied to all protein groups.

### Mouse Behavioral Phenotyping

#### Open field test

Open field testing was conducted as described previously [62]. Briefly, each mouse was placed facing a corner in a 40 cm × 40 cm square-shaped arena and monitored for 10 min to measure locomotor activity. Movements were tracked and analyzed using the AnyMaze software (Stölting).

#### Novel object recognition test (NOR) and Object location test (OLT)

NOR and OLT testing were conducted as described previously [62]. Briefly, each mouse was placed facing a corner in a 40 cm × 40 cm square-shaped arena and monitored for 10 min to measure locomotor activity. Movements were tracked and analyzed using the AnyMaze software (Stölting). Mice were excluded if they failed to explore both objects in either the training or test phase, or if total exploration time in the test phase was less than 10s.

#### Morris water maze

Spatial learning and memory formation were assessed in the Morris water maze [125] as detailed previously [126]. Briefly, the apparatus consisted of a camera, a Perspex platform (10 cm diameter) and a white circular pool (140 cm diameter and 50 cm height) filled with water containing diluted non-irritant white acrylic-based paint dye. The pool was virtually divided into four equal size quadrants, with four different visual cues placed equidistant in the four quadrants. The platform was hidden in the target quadrant (Q1) and submerged 1 cm below the water. The room was low-lit. Testing was done on 6 consecutive days, consisting of three tests. On day 0, a visual cue (visible flag) was placed on top of the escape platform, and four trials were performed. The latency to reach the platform was recorded and mice with an average time greater than 30 s were excluded. On days 1-6, spatial acquisition was performed with each mouse performing four trials starting from different positions in the maze. Individual mice were put into the water facing the wall and allowed 60 s to find the hidden platform where they remained for an additional 60 s. The escape latency was recorded as the time to find the platform. If a mouse failed to reach the platform within 60 s, it was guided to the platform. Probe trials were done on day 6 when each mouse was subjected to a 60 s swim trial without the hidden platform in the maze. The time in each quadrant was recorded. Movements were tracked and analyzed using AnyMaze software (Stölting). Swim traces were classified using a previously described paradigm [127]. Briefly, swim patterns were scored as follows: 1, thigmotaxis; 2, random swim; 3, scanning; 4, chaining; 5, directed search; 6, focal search; and 7, direct swim. Based on this scoring scheme, 1–4 are non-hippocampal strategies, whilst search strategies 5–7 reflect hippocampal learning.

#### Motor tests

Rotarod and pole tests were performed as previously described [128]. Grip strength was performed as previously described [129] using a grip strength meter to measure peak forearm strength (Chatillon, AMETEK). For the beam challenge test, mice were placed at one end of a horizontal pole (47.5 cm from the dowel, 0.8 cm diameter) suspended over a Perspex box, and a nesting house was placed at the far end to motivate the mice to walk across the beam [130]. Mice that failed to complete tasks for non-balance reasons (did not slip or fall) were excluded.

### Knockdown of *Dory* by injection of ASOs into the rat dorsal hippocampus

ASOs targeting rat *Dory* were custom-designed and synthesized by IDT. ASOs incorporated a 5-10-5 Gamper configuration with a combination of phosphorothioate and phosphodiester linkages, as well as 2′-*O*-methoxyethyl modification [131]. The sequences of the oligos are provided in Table 1. Purification of the oligos was performed via HPLC, followed by sodium salt exchange by IDT. Oligos were then ethanol-precipitated prior to formulation. Dry oligos were resuspended in nuclease free water and stored at -20 °C. For *in vivo* delivery, on the day of delivery, oligos were diluted in artificial cerebrospinal fluid (aCSF) prepared with a HEPES buffer solution. The formulation of the aCSF included 137 mM NaCl, 5 mM KCl, 2.3 mM CaCl_2_, 1.3 mM MgCl_2_, 20 mM glucose, and 8 mM HEPES.

**Table 1:**
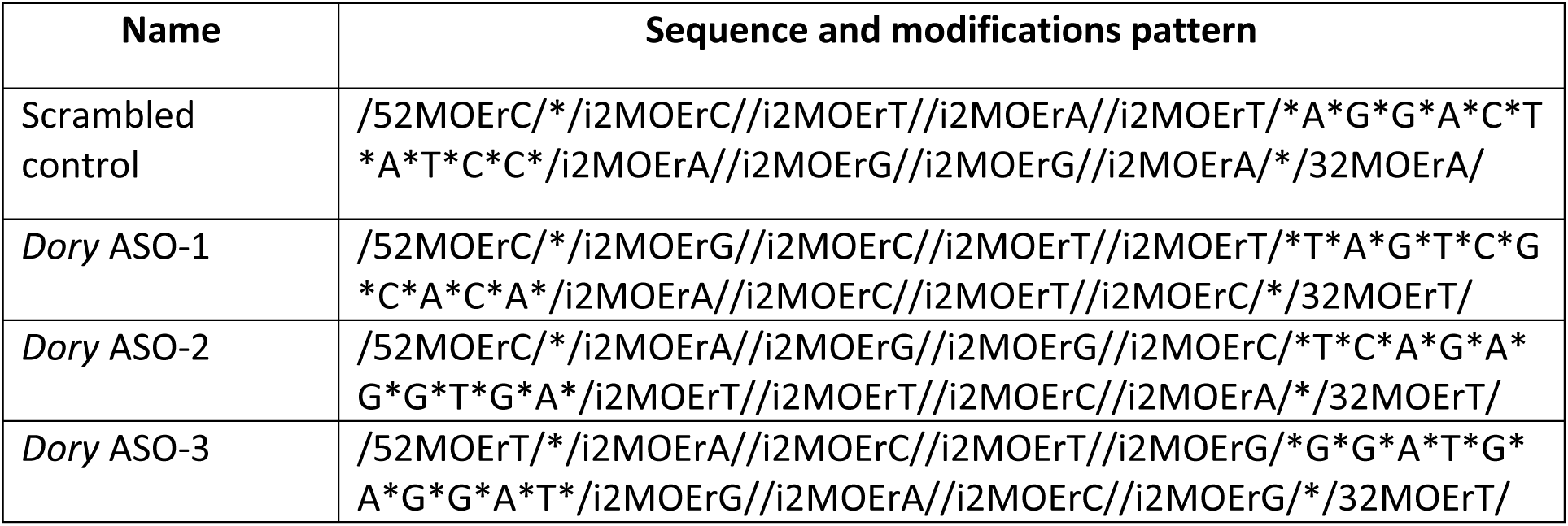
*Dory* antisense oligonucleotide (ASO) sequences. * = phosphorothioate linkage, 2MOEr = 2′-*O*-methoxyethyl on a 5′ (5), internal (i), or 3′ (3) nucleoside. Slashes flank modified nucleosides.

At 2.5-months old, 16 male and 16 female Long-Evans rats were taken for *Dory* knockdown in the dorsal hippocampus. Prior to surgery, rats were handled daily for seven days to acclimatize and reduce stress during the surgery. Rats were anesthetized with isoflurane gas (3-5%) in oxygen (1 L/min) and placed in a stereotaxic frame (David Kopf Instruments). Body temperature was monitored and maintained at 37 °C using a temperature controller. Skin in the surgical area was shaved and cleaned with iodine surgical scrub (Schein, Port Washington, NY). Local anesthesia was administered at the incision site, and a midline incision along the skull was made to expose cranial landmarks (lambda and bregma). The skull was drilled to target the dorsal hippocampus (AP = -3.8, ML = ±2.8, DV = -3.5).

ASOs were infused at a rate of 0.25 μL/min into the dorsal hippocampus, with a dose of 1 µg/µL, 1 µL injected per side. This dose was selected to ensure effective tissue penetration while minimizing toxicity. Given the larger CNS and CSF volume in rats, this direct hippocampal delivery ensured that the ASO reached sufficient concentration at the site of action, in line with previous *in vivo* studies targeting similar brain regions [132–134]. The chosen volume of 1 µL per side was selected to avoid exceeding the tissue capacity, ensuring efficient binding and degradation of the target lncRNA while minimizing tissue damage. Rats were monitored for 15-20 min post-surgery, then moved to separate recovery cages for 12 h before being returned to their home cages. Rats were monitored and weighed daily after surgery and given 7 days to recover before behavioral testing.

### Rat Behavioral Phenotyping

Beginning from seven days following surgery, rats were subjected to behavioral testing using the open field test, novel object recognition test (NOR), object location test (OLT), and Barnes maze test. All analyses were automated using DeepLabCut [135] and SimBA [136]. Neural network models were trained by an experimenter blinded to rat condition. A detailed description of the model training parameters, automated analyses and scripts used are available at https://github.com/rown-h/Dory_Behavioural_Analysis.

#### Open field test

To habituate rats to the testing environment and to assess autonomous movements, an open field test was performed. The test was conducted in a designated testing arena consisting of a black Perspex box (H 63 cm x W 58 cm x L 50 cm), situated in a dimly lit room with an overhead camera positioned centrally. During this phase, each rat was individually transported to a preparation area adjacent to the testing arena and allowed to acclimatize for 20 min in a testing room. Subsequently, the animal was placed in the empty testing arena for 10 min and their movements were recorded. The test arena was wiped down with 70% ethanol between animals to eliminate odor cues. Rats were returned to their home cages and left undisturbed for 24 h prior to the next test.

#### Novel object recognition test (NOR) and Object location test (OLT)

On the day following the open field test, rats underwent the NOR. During the familiarization phase, each rat was transported to a preparation area for a 20 min acclimation period, then placed in the testing arena for 10 min in the presence of two identical objects (either two Spam cans or two glass jars). Object type was counterbalanced across animals. The objects and arena were thoroughly cleaned with 70% ethanol between trials. Exploration time directed toward each object was recorded. Following this phase, rats were returned to their home cages for a 1 h retention interval. They were then brought back to the preparation area for another 20 min before being reintroduced to the arena for a 10 min test phase, now containing one familiar and one novel object.

On the day following the NOR, rats underwent the OLT. Preparation and acclimation procedures prior to testing were identical to those described for the NOR. During familiarization, each rat was placed in the testing arena with two identical objects (glass bottles) and allowed to explore for 10 min. Objects were different to those used in the NOR. Following this phase, rats were returned to their home cages for a 1 h retention interval. In the test phase, rats were reintroduced to the arena for 10 min, where one of the two identical objects had been moved to a new location, while the other remained in its original position. The novel object location was counterbalanced across animals and trials to control for side bias. Both the arena and objects were thoroughly cleaned with 70% ethanol between trials.

For NOR and OLT, exploratory behavior was defined as active investigation of an object, characterized by the rat directing its nose toward the object within a maximum distance of 2 cm, including sniffing, whisker contact, and oral interactions such as licking or biting. Climbing on and resting against the object were not considered exploration. The discrimination index (DI), calculated as the time spent exploring the relocated object relative to total exploration time, provided a measure of hippocampus-dependent spatial memory. A positive DI indicates a preference for the novel object or location, whereas a value near zero suggests equal time spent with both, and a negative value reflects a preference for the familiar item.

The discrimination index was calculated as:

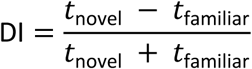

Total exploration time in both training and test phases was calculated. Rats were excluded from analysis if they failed to explore both objects in either the familiarization or test phase, or if total exploration time in the test phase was less than 10s, consistent with previously established criteria [137].

#### Barnes maze test

Prior to testing, the experimental room was prepared to ensure a controlled environment. Illumination was provided by white light (800–1000 lux) which acts as mild aversive stimuli to motivate the rats for searching the shelter in the escape box. All training and test sessions were performed between 8:00 a.m. to 5:00 p.m. A video camera mounted above the Barnes maze arena recorded all movements. Three distinct visible cues were placed evenly on the surrounding curtain, approximately 15 cm above the platform surface, with none positioned directly behind the escape hole. To prevent confounding olfactory influences, no sources of odor (e.g., waste bins) were present in the room. The Barnes maze used in this study was a Columbus Instruments rat maze (model 080344, steel, black, circular, diameter 122 cm, height 80 cm). The platform contained 18 evenly spaced holes, each 9 cm in diameter, through which rats could access the escape box.

Twenty-four hours before the acquisition phase, rats were habituated to the Barnes maze to reduce anxiety-related behavior. Each rat was brought into the testing room and allowed to habituate for 30 min, after which it was placed on the platform and allowed to freely explore for 2 min before being gently guided into the escape box. The escape box was then covered, and the rat remained inside for 1 min before being returned to its home cage. Habituation was conducted under 200 lux illumination without visual cues.

Acquisition training was conducted over five consecutive days, with two training sessions per day separated by a 1 h interval. Each session consisted of a single trial lasting a maximum of 3 min. At the beginning of each trial, the rat was removed from its home cage, brought into the testing room, and allowed to habituate for 5 min. It was then placed at the center of the platform and covered for 15 s with a dark bucket. Upon removal of the bucket, the rat was allowed to explore the maze freely until it located the escape box or until 3 min had elapsed. When a rat entered the escape box, it remained inside for 30 s before being returned to its home cage. If a rat failed to locate the escape box within 3 min, it was gently guided into the box and allowed to remain there for 30 s. To prevent reliance on olfactory cues, the maze surface and escape box were cleaned with 70% ethanol and allowed to air-dry between trials.

A probe trial was performed 24 hours after the final acquisition session (day 6) to assess spatial memory retention. For this trial, the escape box was removed. Each rat was placed under the start box for 15 s, after which the start box was lifted, and the animal was allowed to explore the maze freely for 90 s. During this period, primary latency and primary errors to the escape hole location were recorded.

Numerous parameters were measured to assess spatial learning and memory in the Barnes maze test. Primary latency was defined as the time elapsed between release from the start box to the first head poke into the escape box. Total latency measured the time from release to complete body entry into the escape box. Similarly, primary and total path length quantify the distance travelled a rat’s center before these respective timepoints, and primary and total errors quantify the number of incorrect holes searched before the first head poke or complete entry into the escape box respectively. Finally, search strategies were scored using an 11-point system [138] based on the animal’s search pattern before first head poke: direct (11), corrections within escape box sector (9-10), corrections outside escape box sector (7-8), serial search patterns (2-8), and random search (1). These measures accounted for instances in which animals located the escape box but delayed entry due to exploratory behavior.

### Statistical analysis

Statistical analysis was performed in GraphPad Prism version 10. Statistical significance was determined using unpaired t-tests for two-group experiments. One-way analysis of variance (ANOVA) tests were conducted to compare more than two datasets, and two- and three-way ANOVA tests were applied to compare groups across time. All behavioral and imaging quantification data are presented as mean ± standard error (SEM). Molecular data are presented as mean ± standard deviation (SD). Details on individual test parameters and group numbers are provided in figure legends.

## Data Availability Statement

RNA capture sequencing reads area available from the Sequencing Read Archive (SRA) under PRJNA1417931. RNA sequencing reads from *Dory* wt/wt and Δ/Δ hippocampus are available under PRJNA1417287. Mass spectrometry data (.raw files) from *Dory* wt/wt and Δ/Δ mice hippocampi are available on Zenodo (https://doi.org/10.5281/zenodo.18607396).

## Acknowledgements

This study was supported by grants DP210103233 and DP210101957 from the Australian Research Council. J.S.M. and M.C. are supported by SHARP Fellowship RG193211 from UNSW Sydney. We thank Jessica Spathos, Andrea Chan and Annika van Hummel (Macquarie University) and Lachlan Ferguson (UNSW) for technical assistance, Simon Nilsson (University of Washington) for assistance with SimBA video analysis software, Kelly Clemens (UNSW) for access to rat RNA sequencing data for ASO design, and Mike Clark (University of Melbourne) for assistance with RNA capture sequencing.

## Author contributions

J.S.M., F.D. and L.M.I. designed the study and oversaw the work. F.D. and S.J. designed and generated the knockout mice. S.J. and M.C. performed and analyzed the Sanger sequencing. M.C. planned and performed the RNA sequencing and RNA capture sequencing transcript isoform identification. S.J. and S.A. performed the *in situ* hybridization. A.P. performed and analyzed mass spectrometry data. S.J. performed mass spectrometry analyses and all mouse behavioral tests. K.C., L.F. and S.A. designed and performed rat knockdown experiments. R.J.H. analyzed rat knockdown behavior tests. S.J., M.C. and J.S.M drafted the manuscript, which was refined and edited by all authors.

## Supplemental information

Document S1. Supplementary Figs. S1-S6.

Supplementary File S1 Differentially expressed genes and proteins.

Supplementary File S2 RNA sequencing.

Supplementary File S3 Mass spectrometry.

## Supplemental information

**Fig. S1:**
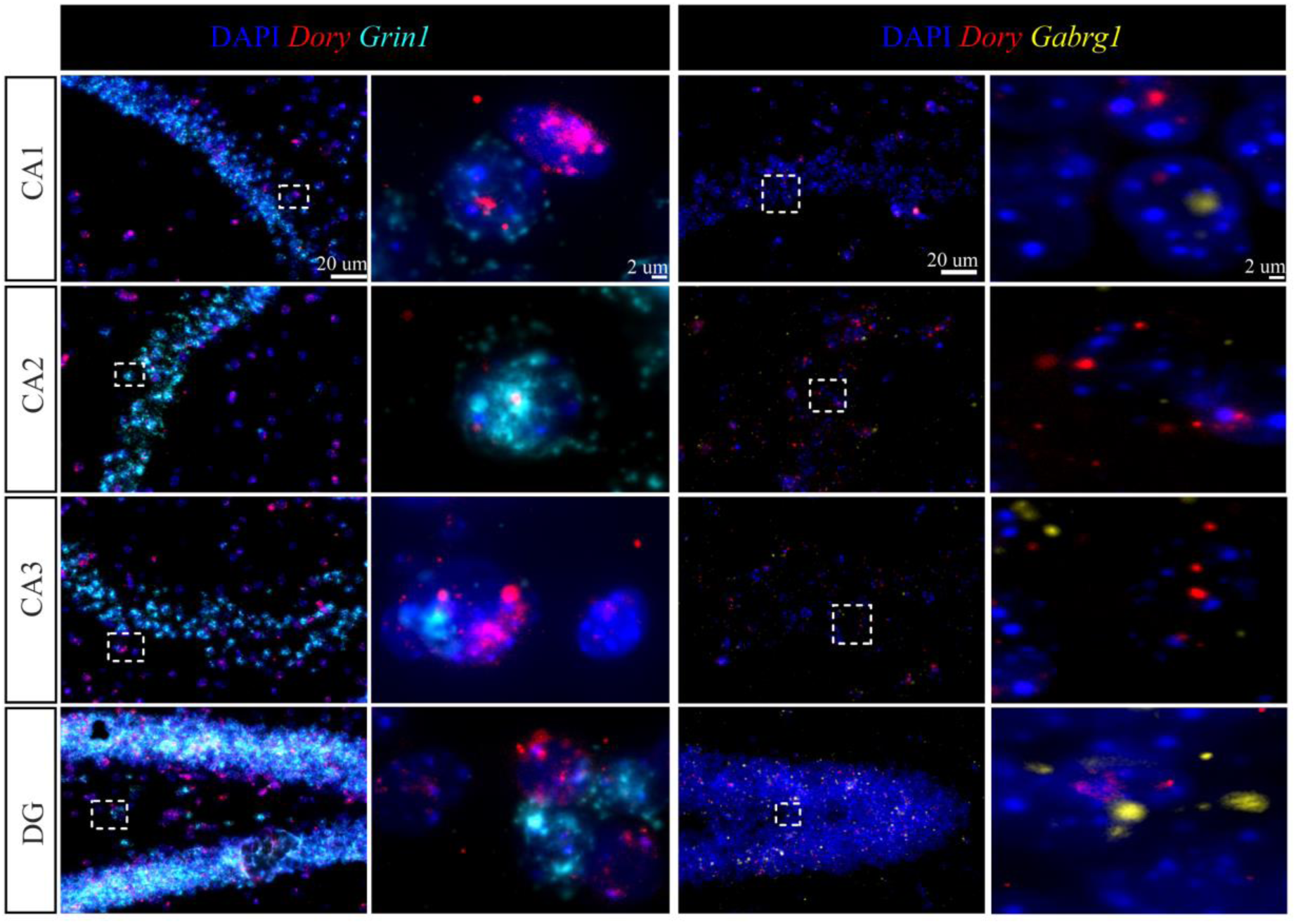
Representative images of Dory labelling in mouse hippocampus. Hippocampus containing brain sections were labelled with DAPI, *Dory* and *Grin1* for RNAscope colocalization. CA – *cornu ammonis* (with subfields 1–3 shown), DG – dentate gyrus.

**Fig. S2:**
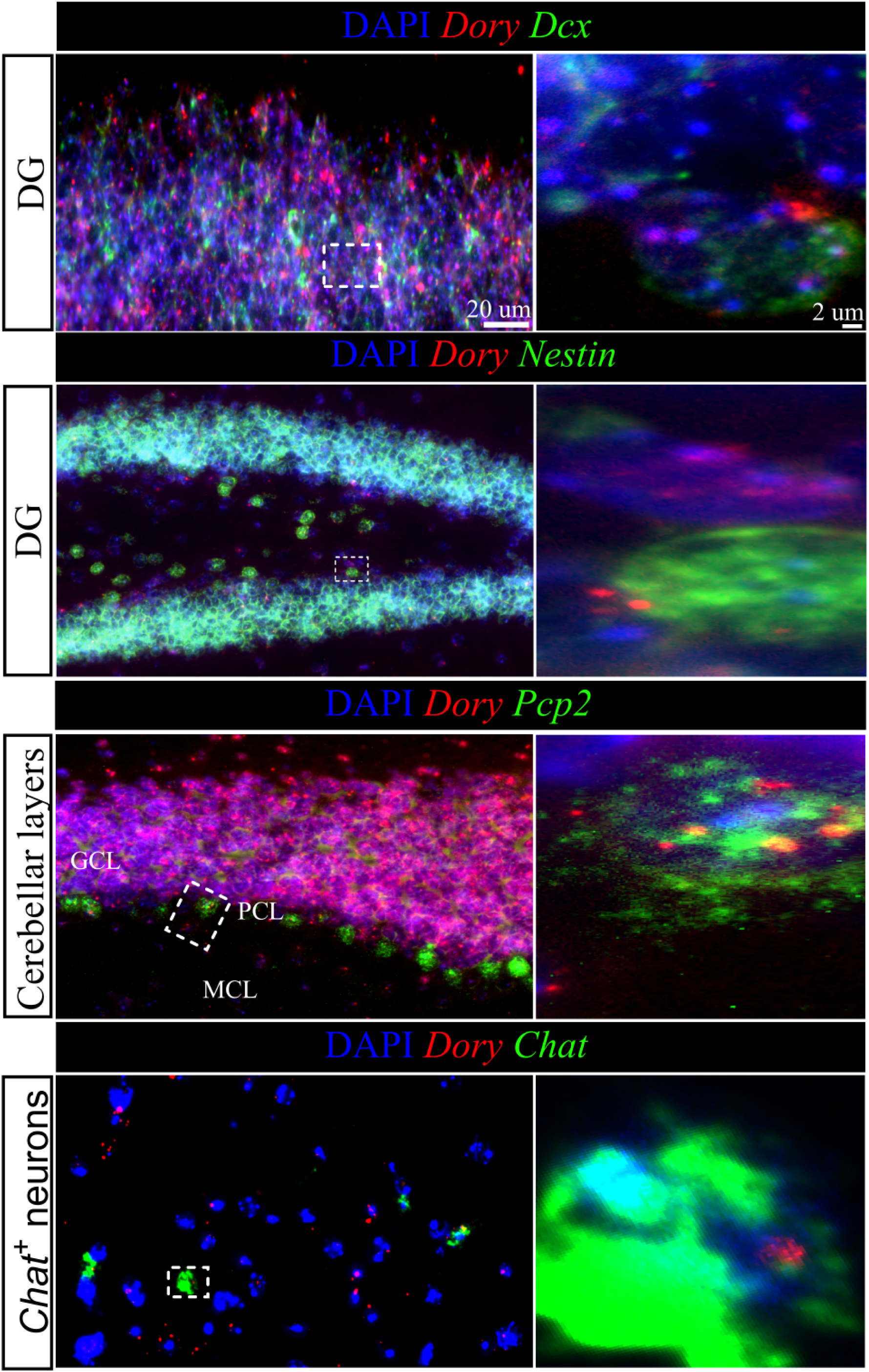
Representative images of *Dory* labelling in mouse brain. Brain sections were labelled with DAPI, *Dory* and neuronal subtype markers (Dcx – immature neurons; Nestin – neural stem cells; Pcp2 – Purkinje cells; Chat – cholinergic cells) for RNAscope colocalization. GCL – granular cell layer; PCL – Purkinje cells layer; MCL – molecular cell layer. DG – dentate gyrus.

**Fig. S3:**
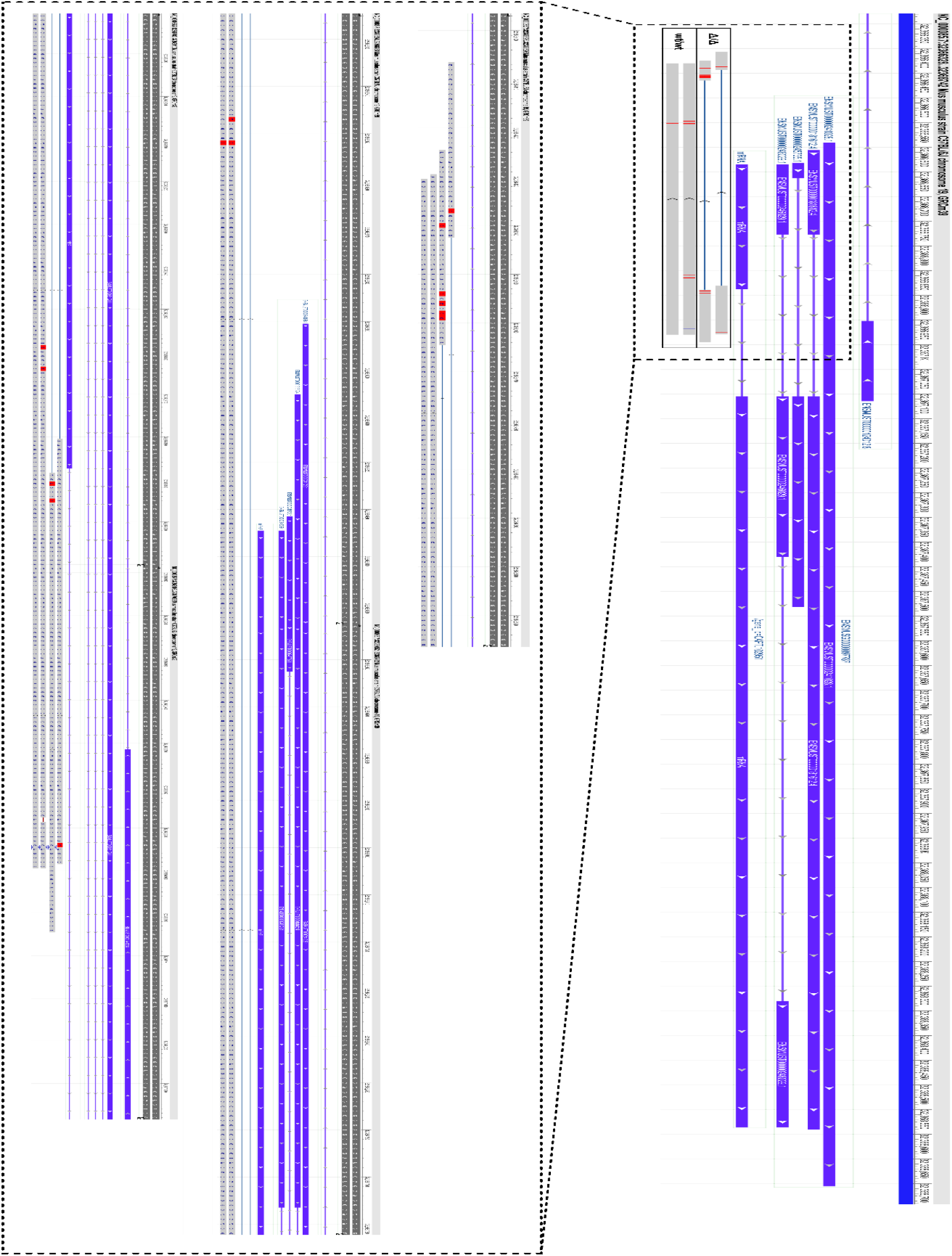
Sanger sequencing results. Amplicons from F2 generation mice were sequenced using Sanger sequencing. Amplicons were aligned to the GRCm39 mouse genome using NCBI blastn. Low quality bases were trimmed from the end of amplicons.

**Fig. S4:**
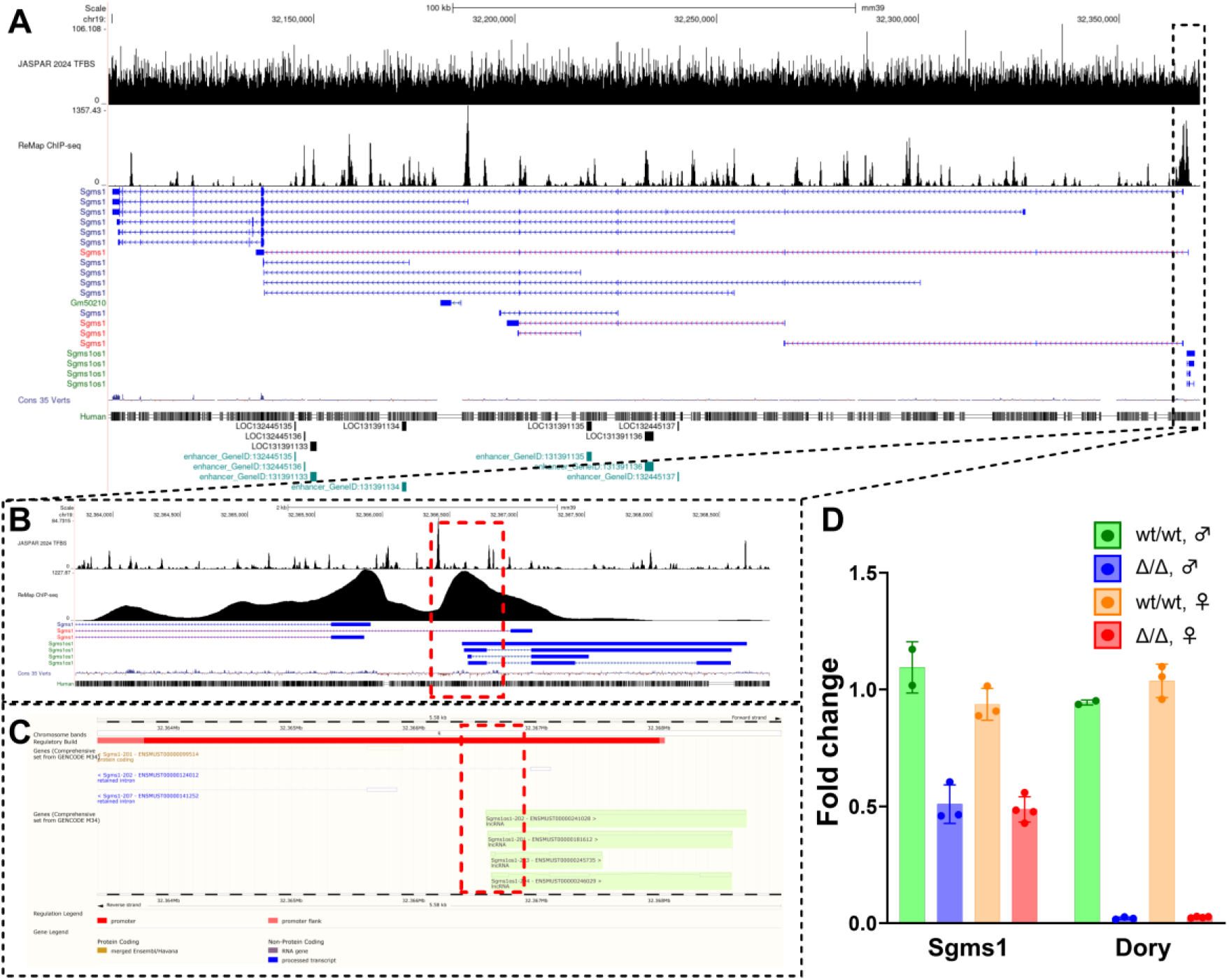
Detailed view of the *Dory* (*Sgms1os1/2700046G09Rik*) knockout locus. (**A**) UCSC track browser view of the *Sgms1* and *Sgms1os1* (*2700046G09Rik*) locus on mouse chr19, detailing density of transcription factor binding sites (JASPAR TFBS), ReMap CHIP-seq peaks, transcripts at the locus, base conservation (PhyloP), human multi-alignment, and enhancer loci. (**B**) Zoomed-in view of the knockout locus (red box). ReMap CHIP-seq density shows 2 peaks associated with the transcription start sites (TSS) of *Sgms1* and *Sgms1os1* (*2700046G09Rik*). The deletion only covers the latter. (C) Ensembl genome browser showing the presence of a promoter (red bar) at the knockout locus (red box). (D) RNA-seq fold changes of Sgms1 and Sgms1os1 in knockout mice.

**Fig. S5:**
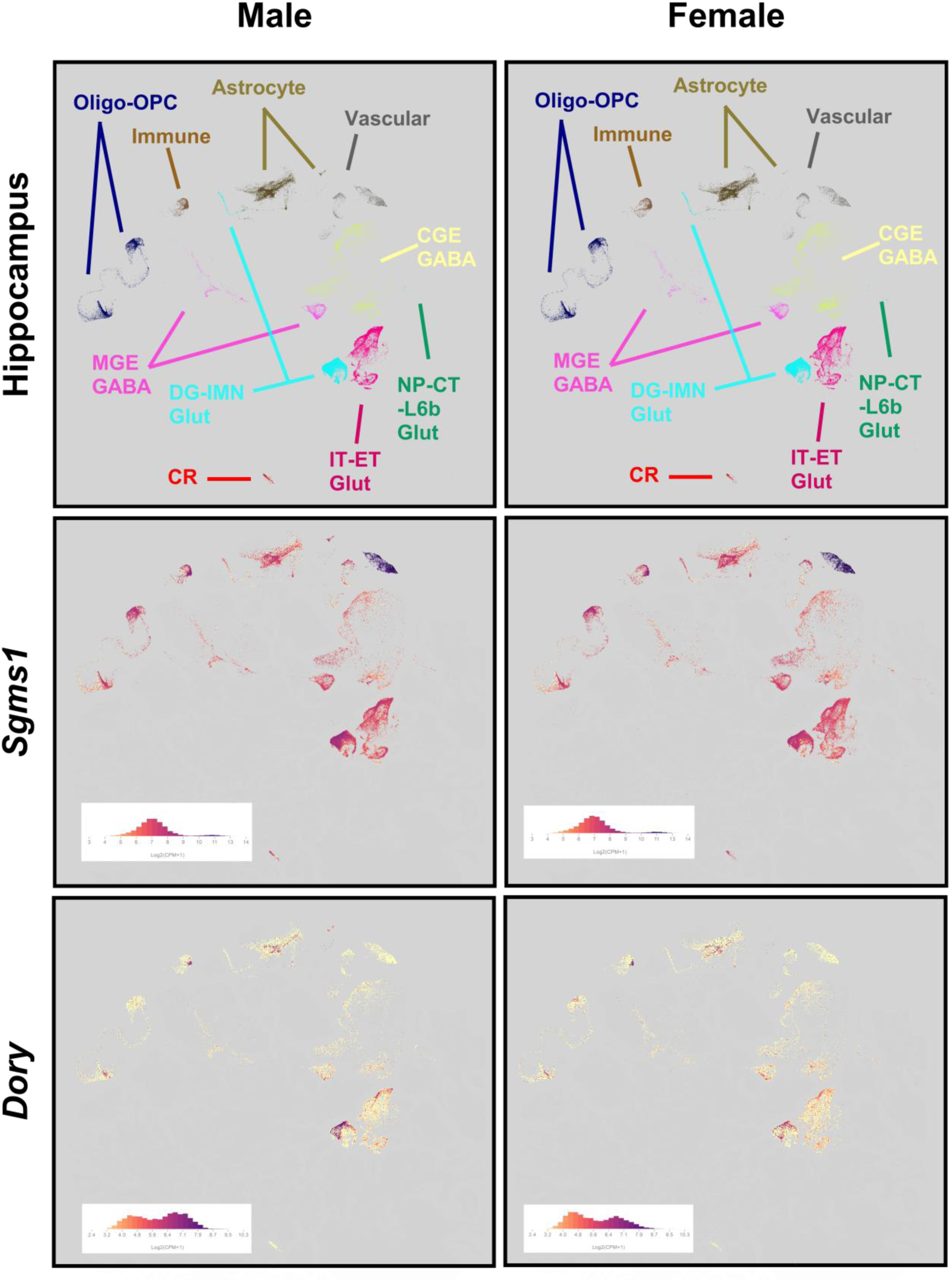
*Sgms1* is ubiquitously expressed in the hippocampus while *Dory* is enriched in neurons. UMAPs of Allen Brain Cell Atlas hippocampus male and female cells (RRID:SCR_024440; https://portal.brain-map.org/atlases-and-data/bkp/abc-atlas10.1038/s41586-023-06812-z). For hippocampus panels, cells are colored by cell cluster. For Sgms1 and Dory panels cells are colored by RNA-seq expression value (log(CPM+1)). CGE - Caudal Ganglionic Eminence, CPM - counts per million, CR - Cajal-Retzius, CT - Corticothalamic, DG - Dentate Gyrus, ET - Extratelencephalic, GABA - Gamma-Aminobutyric Acid, Glut - Glutamate, IMN - Immature Neuron, IT - Intratelencephalic, L6b - Layer 6b, MGE - Medial Ganglionic Eminence, NP - Near-Projecting, Oligo - Oligodendrocyte, OPC - Oligodendrocyte Progenitor Cells.

**Fig. S6:**
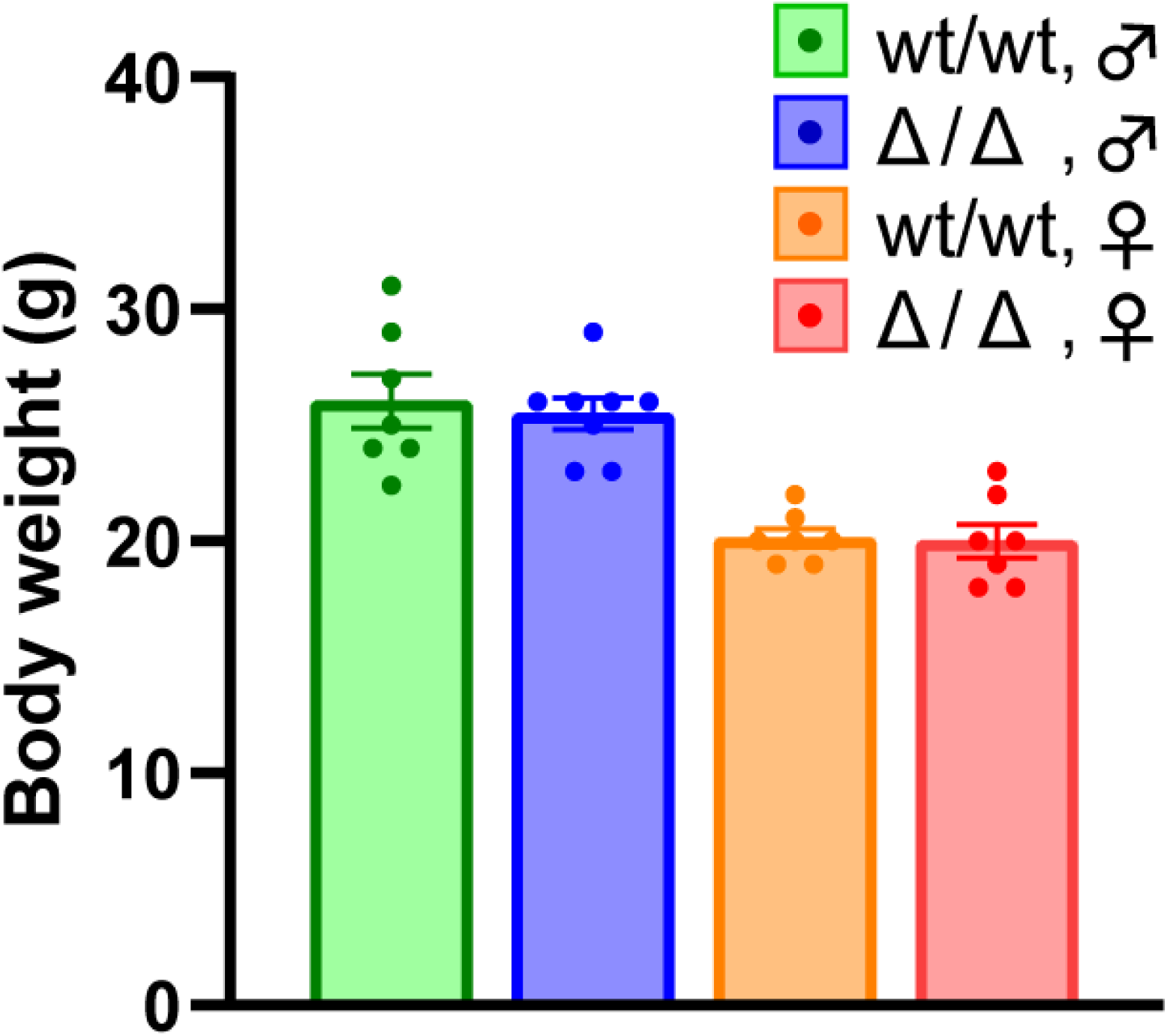
Body weights of wild-type and Dory knockout mice. There were no significant differences by unpaired t-test in weights between controls and *Dory* Δ/Δ mice in either sex (n=7-8/group).

**Fig. S7:**
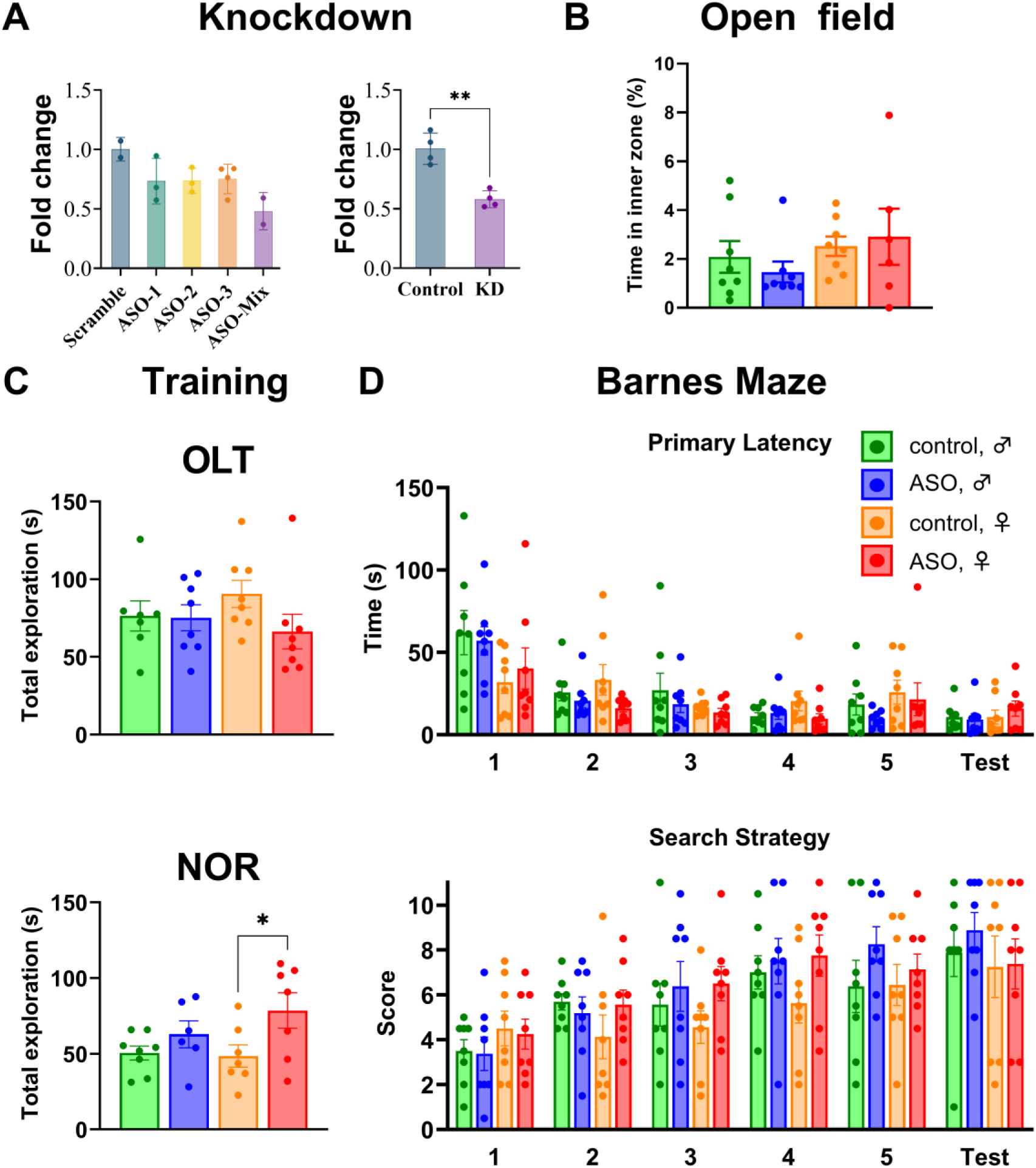
Preliminary knockdown of Dory by antisense oligonucleotides (ASOs) and extended rat knockdown behavioral results. (A) Preliminary test by qPCR of (left) different ASOs for Dory knockdown (n=2-4/group) and (right) knockdown of *Dory* using ASO mix after 1 week post-surgery in dorsal hippocampus. (n=4/group). (B) No differences in open field time in center (n=6-8/group). (C) Training total exploration time for the object location task (OLT) and novel object recognition task (NOR). (D) Barnes maze results in Dory knockdown rats (n=8/group). We did not find any differences between controls and knockdown rats in primary latency over training days or on the test day. We found a significant (p=0.0355) increase in search strategy score in knockdown rats (5.7 vs 6.5), however the effect size was smaller than score measure.

**Fig. S8:**
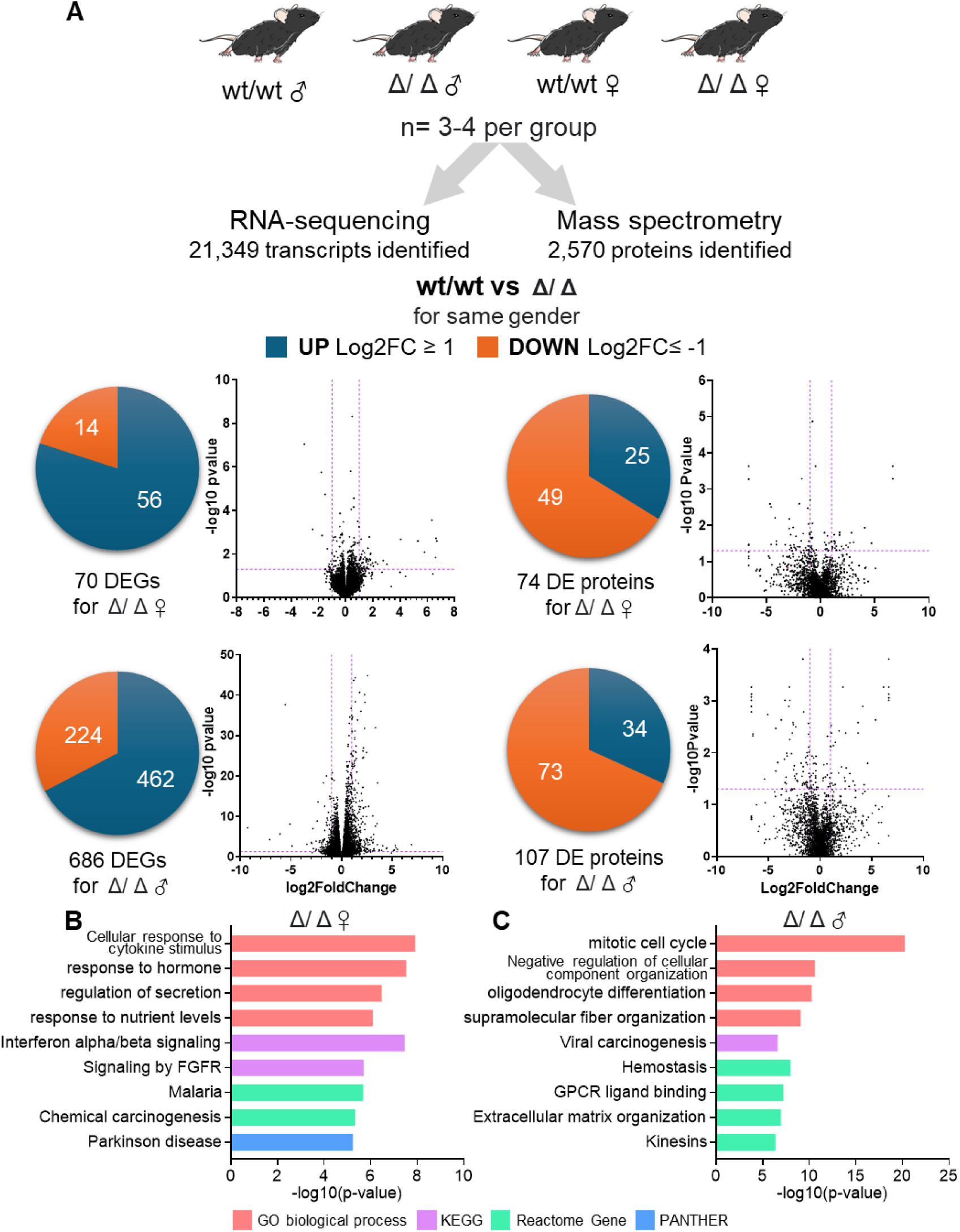
Analysis of RNA-sequencing and mass spectrometry data from the hippocampus of *Dory* knockout mice. (A) Left, RNA sequencing (21,349 genes quantified). Right, mass spectrometry. (2,570 proteins quantified). The number of differentially expressed genes (DEGs) and differentially expressed proteins (DEprots) with mean log2FC ≥ 1 or ≤ -1, and p-value < 0.05 are specified. Presented are DEGs and DEPs volcano plots, respectively. (B) Male and (C) female *Dory* Δ/Δ GO enrichment, KEGG pathway, Reactome gene and PANTHER analysis of hippocampus DEGs and DEProts.

