## Supplemental Figures for "The long noncoding RNA *Dory* is required for female but not male spatial learning and memory"

### Supplemental information

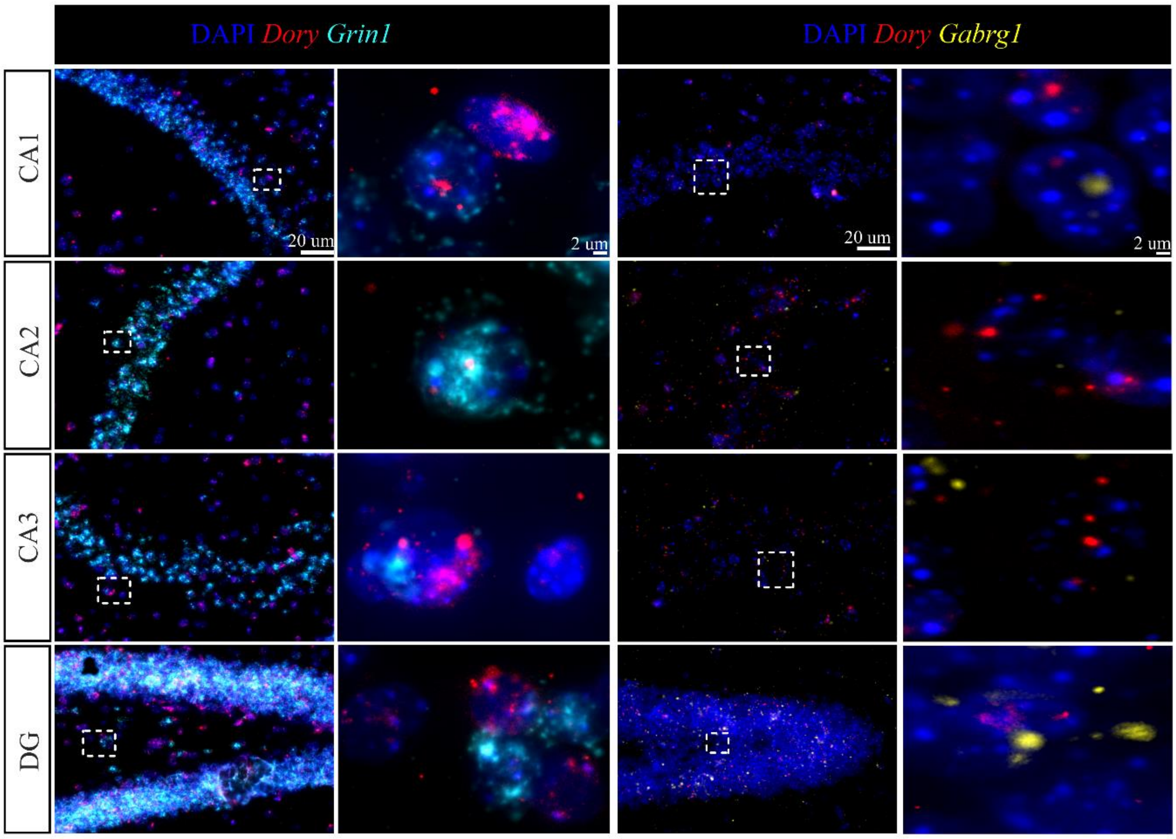


**Fig. S1: Representative images of *Dory* labelling in mouse hippocampus.** Hippocampus containing brain sections were labelled with DAPI, *Dory* and *Grin1* for RNAscope co-localization. DG – dentate gyrus.

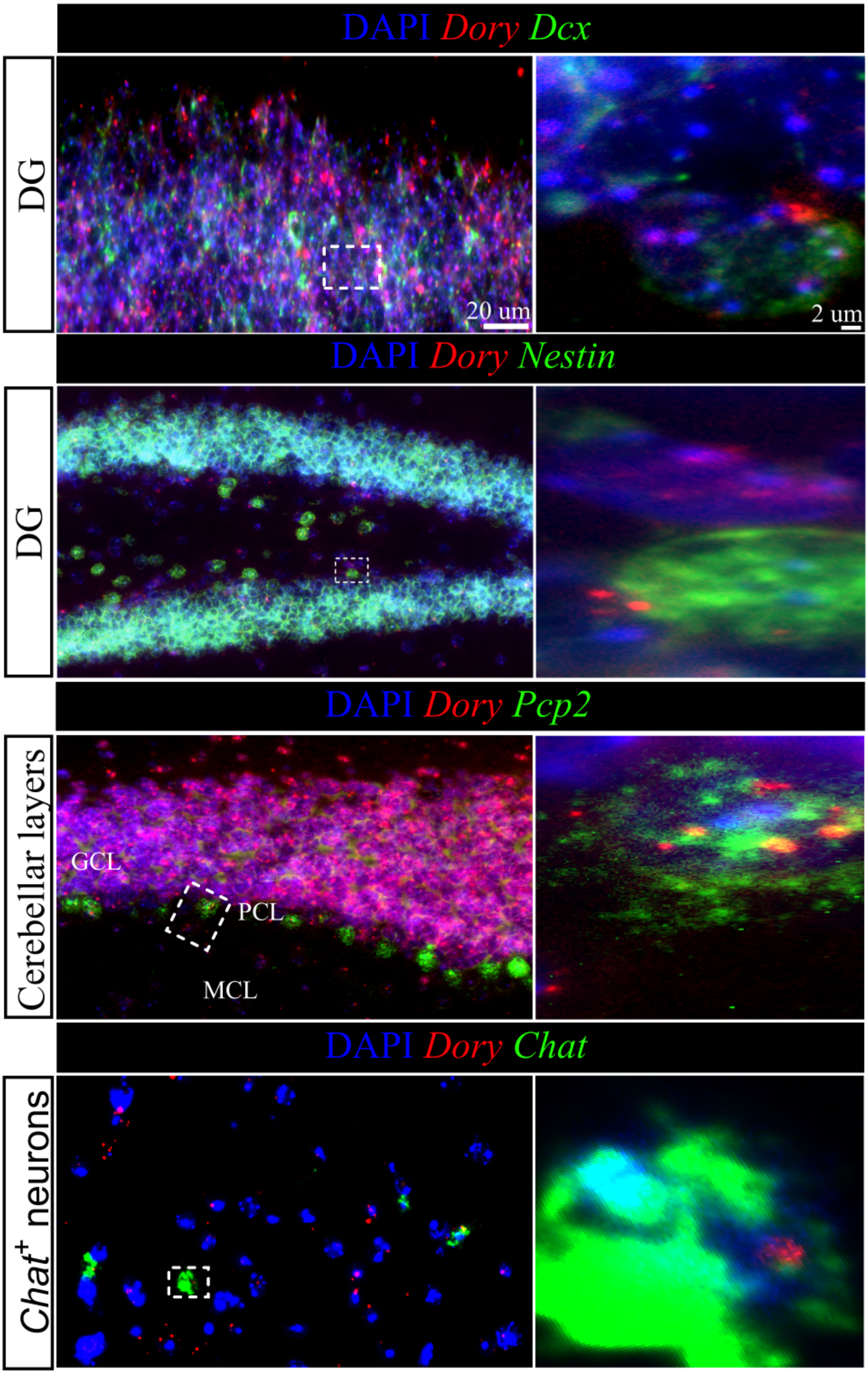
DAPI Dory Dcx
DG
20 um
2 um
DAPI Dory Nestin
DG
DAPI Dory Pcp2
Cerebellar layers
GCL
PCL
MCL
DAPI Dory Chat
Chat+ neurons

**Fig. S2: Representative images of *Dory* labelling in mouse brain.** Brain sections were labelled with DAPI, *Dory* and neuronal subtype markers (Dcx – immature neurons; Nestin – neural stem cells; Pcp2 – Purkinje cells; Chat – cholinergic cells) for RNAscope co-localization. GCL – granular cell layer; PCL – Purkinje cells layer ; MCL – molecular cell layer. DG – dentate gyrus.

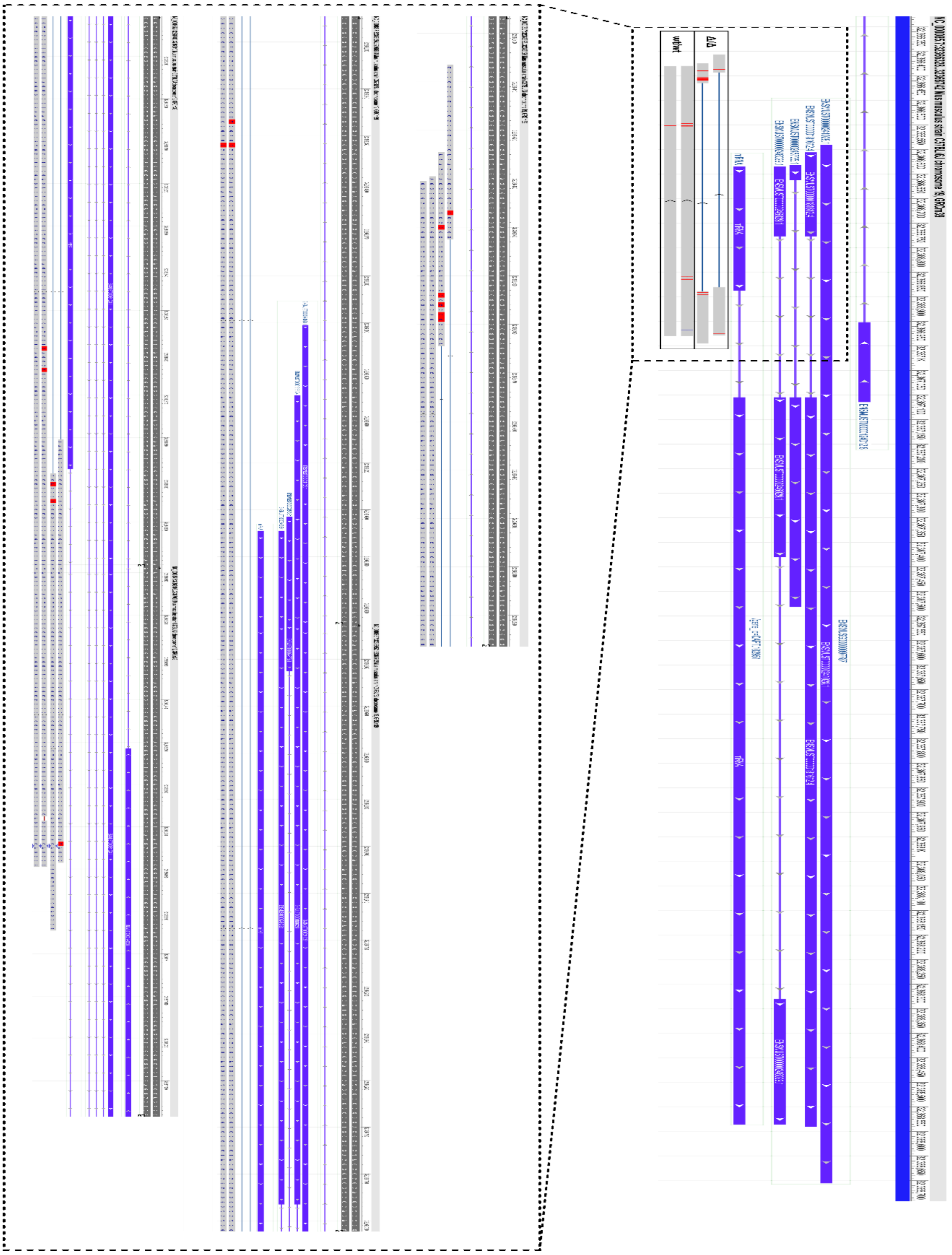

**Fig. S3: Sanger sequencing results**. Amplicons from F2 generation mice were sequenced using Sanger sequencing. Amplicons were aligned to the GRCm39 mouse genome using NCBI blastn. Low quality bases were trimmed from the end of amplicons.

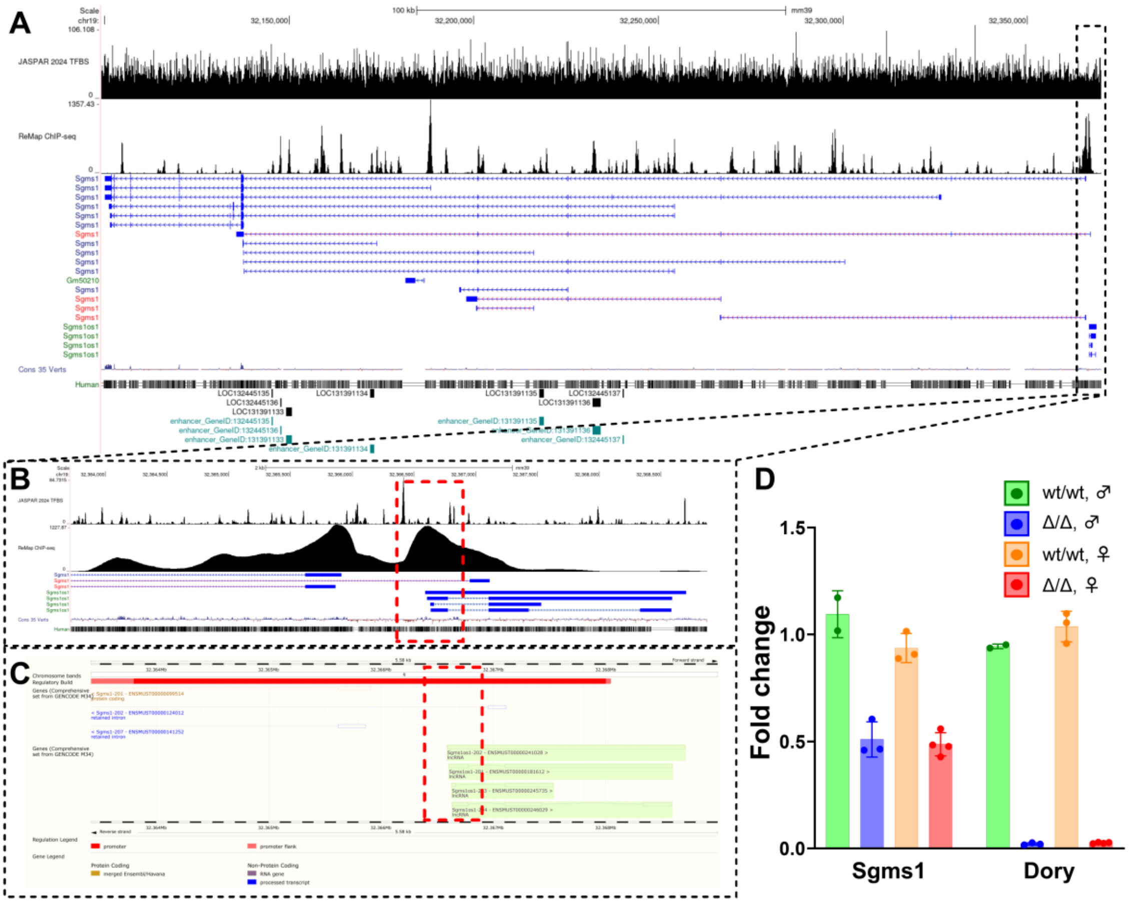


**Fig. S4: Detailed view of the *Dory* (*Sgms1os1/2700046G09Rik*) knockout locus. (A)** UCSC track browser view of the *Sgms1* and *Sgms1os1* (*2700046G09Rik*) locus on mouse chr19, detailing density of transcription factor binding sites (JASPAR TFBS), ReMap CHIP-seq peaks, transcripts at the locus, base conservation (PhyloP), human multi-alignment, and enhancer loci. **(B)** Zoomed in view of the knockout locus (red box). ReMap CHIP-seq density shows 2 peaks associated with the transcription start sites (TSS) of *Sgms1* and *Sgms1os1* (*2700046G09Rik*). The deletion only covers the latter. **(C)** Ensembl genome browser showing the presence of a promoter (red bar) at the knockout locus (red box). **(D)** RNA-seq fold changes of Sgms1 and Sgms1os1 in knockout mice.

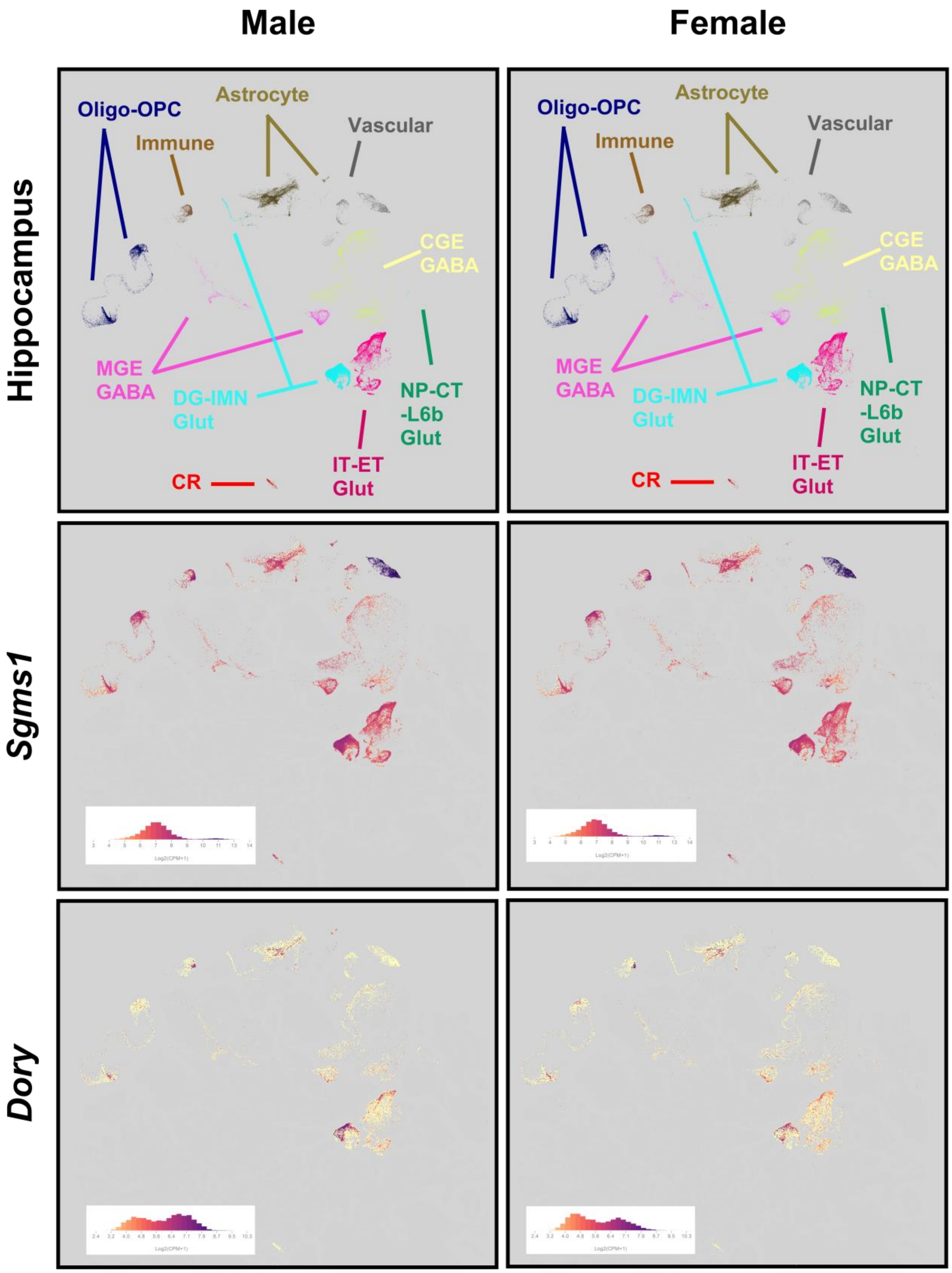
Male
Female
Hippocampus
Sgms1
Dory
Oligo-OPC
Astrocyte
Immune
Vascular
CGE GABA
MGE GABA
DG-IMN Glut
NP-CT -L6b Glut
IT-ET Glut
CR

**Fig. S5: *Sgms1* is ubiquitously expressed in the hippocampus while *Dory* is enriched in neurons**. UMAPs of Allen Brain Cell Atlas hippocampus male and female cells (RRID:SCR_024440; https://portal.brain-map.org/atlases-and-data/bkp/abc-atlas 10.1038/s41586-023-06812-z). For hippocampus panels cells are coloured by cell cluster. For Sgms1 and Dory panels cells are coloured by RNA-seq expression value (log(CPM+1)). CGE - Caudal Ganglionic Eminence, CPM - counts per million, CR - Cajal-Retzius, CT - Corticothalamic, DG - Dentate Gyrus, ET - Extratelencephalic, GABA - Gamma-Aminobutyric Acid, Glut - Glutamate, IMN - Immature Neuron, IT - Intratelencephalic, L6b - Layer 6b, MGE - Medial Ganglionic Eminence, NP - Near-Projecting, Oligo - Oligodendrocyte, OPC - Oligodendrocyte Progenitor Cells.

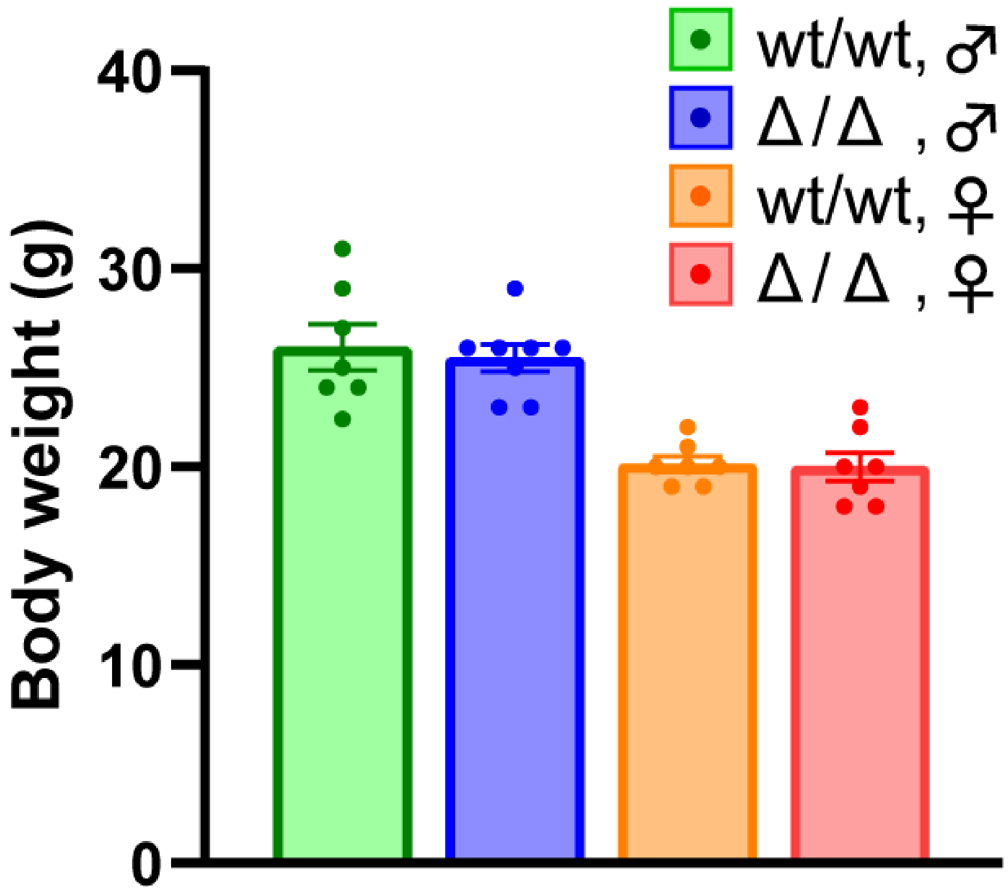


**Fig. S6: Body weights of wild-type and *Dory* knockout mice.** There were no significant differences by unpaired t-test in weights between controls and *Dory* Δ/Δ mice in either sex (n=7-8/group).

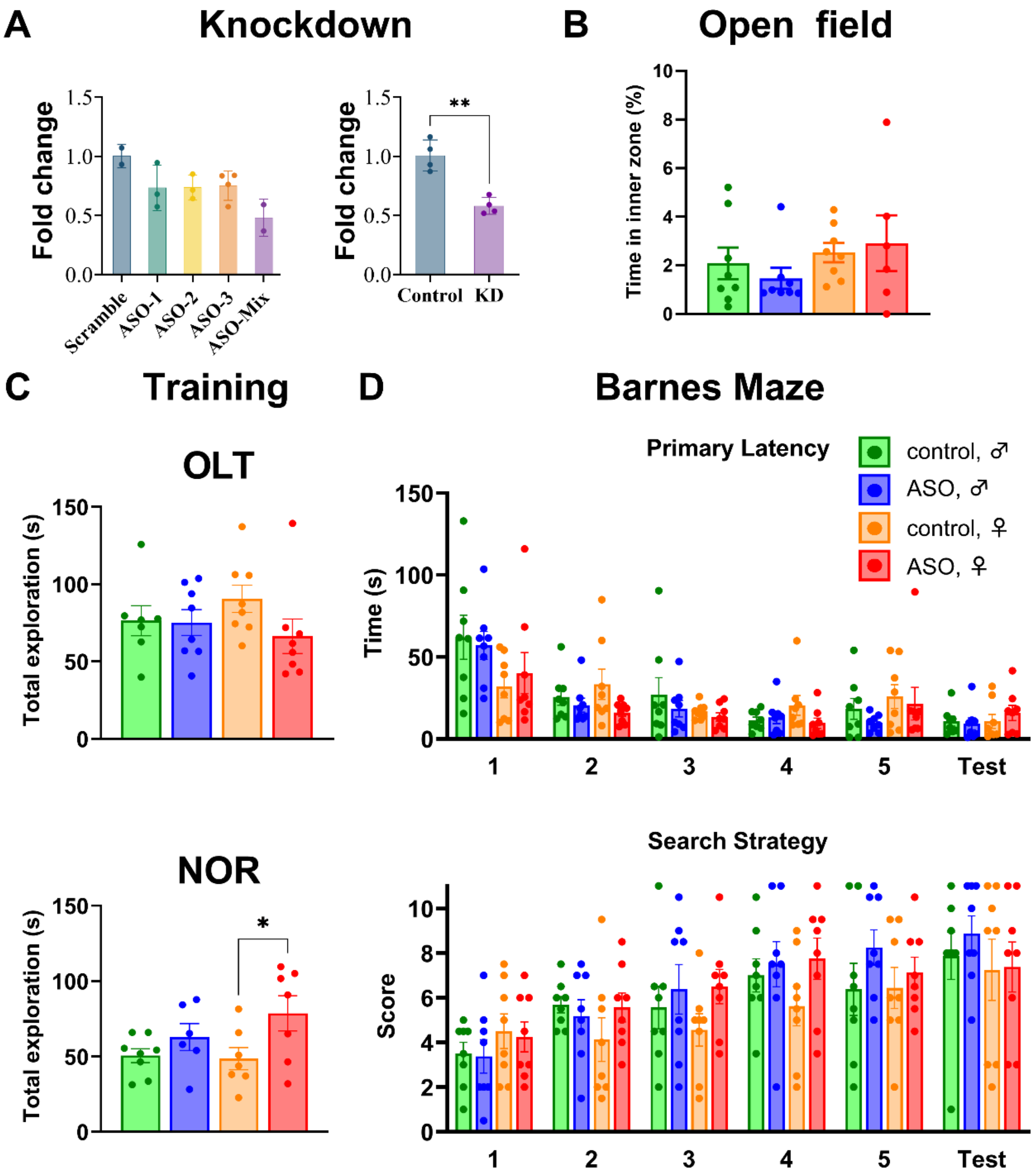


**Fig. S7: Preliminary knockdown of *Dory* by antisense oligonucleotides (ASOs) and extended rat knockdown behavioral results. (A)** Preliminary test by qPCR of (left) different ASOs for *Dory* knockdown (n=2-4/group) and (right) knockdown of *Dory* using ASO mix after 1 week post-surgery in dorsal hippocampus. (n=4/group). **(B)** No differences in open field time in center (n=6-8/group). **(C)** Training total exploration time for the object location task (OLT) and novel object recognition task (NOR). **(D)** Barnes maze results in Dory knockdown rats (n=8/group). We did not find any differences between controls and knockdown rats in primary latency over training days or on the test day. We found a significant (p=0.0355)

increase in search strategy score in knockdown rats (5.7 vs 6.5), however the effect size was smaller than score measure.

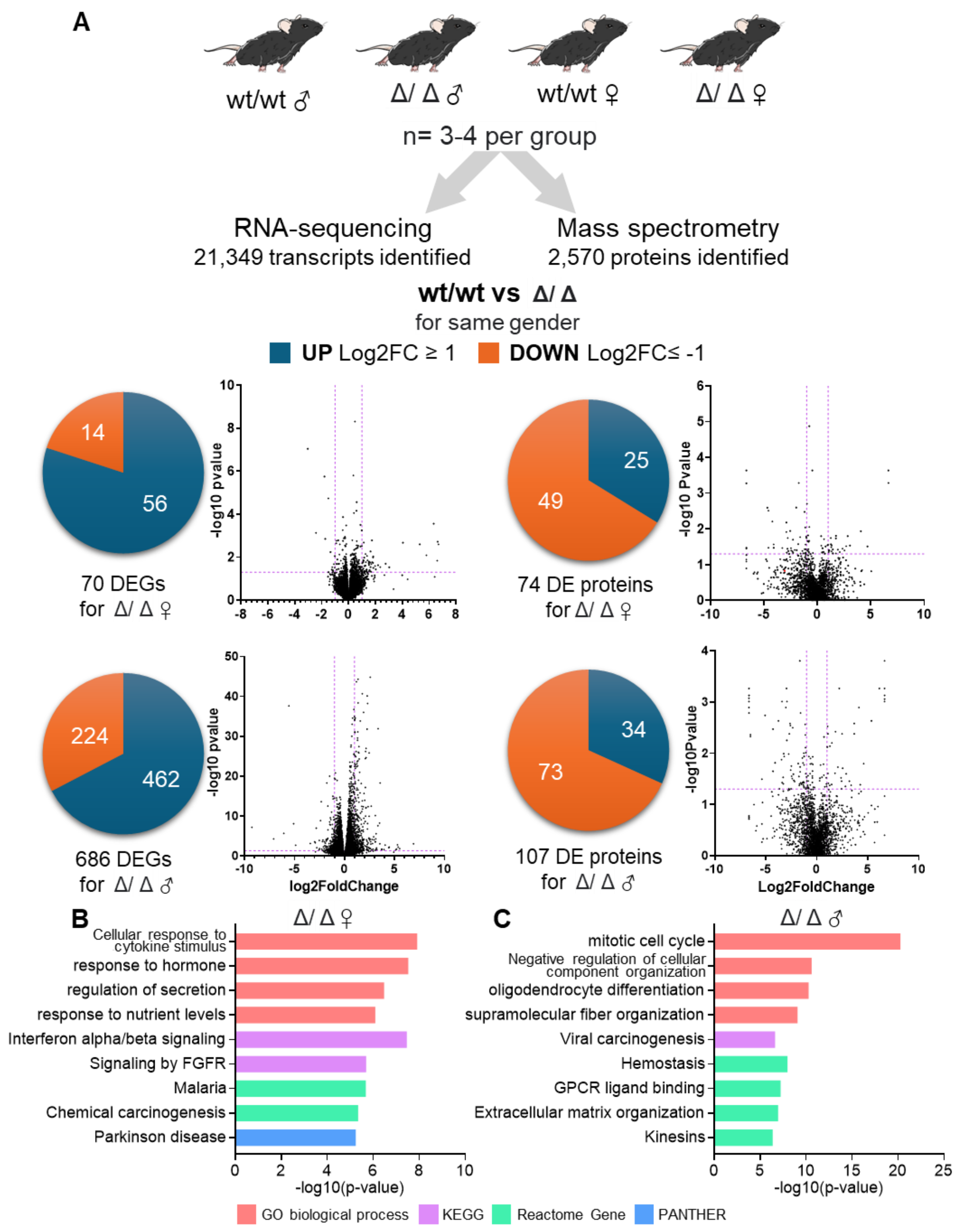


**Fig. S8: Analysis of RNA-sequencing and mass spectrometry data from the hippocampus of *Dory* knockout mice. (A)** Left, RNA sequencing (21,349 genes quantified). Right, mass spectrometry. (2,570 proteins quantified). The number of differentially expressed genes (DEGs) and differentially expressed proteins (DEprots) with mean $log_2FC \geq 1$ or $\leq -1$, and p-value < 0.05 are specified. Presented are DEGs and DEPs volcano plots, respectively. **(B)** Male and **(C)** female *Dory* Δ/Δ GO enrichment, KEGG pathway, Reactome gene and PANTHER analysis of hippocampus DEGs and DEProts.
